# Neural circuits mediating visual stabilization during active motion in zebrafish

**DOI:** 10.1101/566760

**Authors:** Sha Sun, Zhentao Zuo, Michelle Manxiu Ma, Chencan Qian, Lin Chen, Wu Zhou, Kim Ryun Drasbek, Liu Zuxiang

## Abstract

Visual stabilization is an inevitable requirement for animals during active motion interaction with the environment. Visual motion cues of the surroundings or induced by self-generated behaviors are perceived then trigger proper motor responses mediated by neural representations conceptualized as the internal model: one part of it predicts the consequences of sensory dynamics as a forward model, another part generates proper motor control as a reverse model. However, the neural circuits between the two models remain mostly unknown. Here, we demonstrate that an internal component, the efference copy, coordinated the two models in a push-pull manner by generating extra reset saccades during active motion processing in larval zebrafish. Calcium imaging indicated that the saccade preparation circuit is enhanced while the velocity integration circuit is inhibited during the interaction, balancing the internal representations from both directions. This is the first model of efference copy on visual stabilization beyond the sensorimotor stage.

## INTRODUCTION

Accurate perception, especially a keen visual perception, is a significant challenging behavioral requirement for prey capturing, escaping and mating. However, all visually guided animals are faced with retinal image degradation caused by self-generated body motion (Cullen, 2004). To maintain a stable vision during locomotion, many reflexes, such as vestibulo-ocular reflex (VOR), optokinetic reflex (OKR) and proprioceptive reflexes, are required for minimizing retinal slip via fine adjustments of the eye/head in vertebrates (Angelaki and Hess, 2005), known as active visual stabilization. During the past three decades, researches have scrutinized into the mechanism of active visual stabilization by taking different animal models into consideration, e.g. mice (Andreescu et al., 2005), rats (Yoder et al., 2011), cats (Godaux and Vanderkelen, 1984), monkeys (Knight, 2012), and even turtles (Rosenberg and Ariel, 1996).

One potential mechanism underpinning active visual stabilization is to measure the sensory change induced by eye-head movements and to compensate it by feedback motor controls (Sun and Goldberg, 2016). However, its scope has been limited by the processing speed of the visual system, especially in complex coordinated movements, such as eye-head/body interaction or smooth limb control. Instead, another mechanism named efference copy by von Holst (von Holst E, 1950) or corollary discharge (CD) by Sperry (Sperry, 1950), has been demonstrated to be more feasible for gaze stabilization via body adjustment (Lisberger, 2009; Sommer and Wurtz, 2002, 2008). By sending out a copy of the motor commands (efference copy) that generates a predictive representation, this mechanism modulates self-generated sensory inputs by sensory suppression (Lisberger, 2009) or remapping (Wurtz, 2018). This approach enables a calibrated perceptual model of the environments. In spite of the sensory modulation, the efference copy evokes compensatory eye movements, as a direct motor compensation, especially during rhythmic body movements (Easter and Johns, 1974; Wolpert and Miall, 1996), to minimize the self-generated sensory changes. Recently, one source of this modulation has been identified in the spinal central pattern generator (CPG), which evokes tail undulation in general but also has a fast ascending pathway to control eye movements, even in the absence of visual input (Combes et al., 2008; Stehouwer, 1987). This projection from the CPG to abducens nucleus is believed to underscore the compensatory eye movements directly during locomotion (Lambert et al., 2012), given the fact that the latency of eye-tail synchrony is nearly zero (Chagnaud et al., 2012). However, considering the role of efference copy in this context, one piece of the puzzle is still missing, between the sensory modulation and the motor compensation approaches. It is unclear if and how the two approaches co-operate with each other, due to the fact that a weighting mechanism is necessary when the two happen simultaneously. It is especially interesting to know whether efference copy interacts with the sensory and motor systems at the same time, while the visual environment is constantly changing, thus, the visual system is occupied by active processing. However, during active visual perception, the self-generated movements always lead to locomotion accompanied with unstable head position, which makes neural recording very challenging.

In this study, we utilize the well-established larval zebrafish model system, majorly benefiting from its translucent brain for neuronal level activity recording via advanced imaging methods during visual behaviors. By comparing the OKR eye movements evoked by whole-field rotating gratings between tail-free and tail-immobilized conditions, we found that tail-beats induced extra reset saccades during OKR. Calcium imaging acquired by two-photon microscopy and light-sheet microscopy revealed enhanced activities in rostral hindbrain and suppressed dorsal-caudal hindbrain for tail-free fish. These results together suggest a third approach by which efference copy interacts with internal representations during active visual perception.

## RESULTS

### Tail-eye interactions during OKR observed in behavioral assays

We used a well-established paradigm to elicit OKR (Portugues et al., 2014) in zebrafish larvae that were restrained in low melting agarose (Easter and Nicola, 1997). Agarose was removed from the eyes and tail (Figure 1A). A rotating, whole-field grating stimulus projected on a screen below the fish (Figure 1B), reliably evoked OKR eye movements (Huang and Neuhauss, 2008). Eye and tail movements in response to the gratings were recorded using an infrared camera (Figure 1A) and eye/tail positions were measured offline from each frame of the acquired videos (Figure 1B).

**Figure 1.**
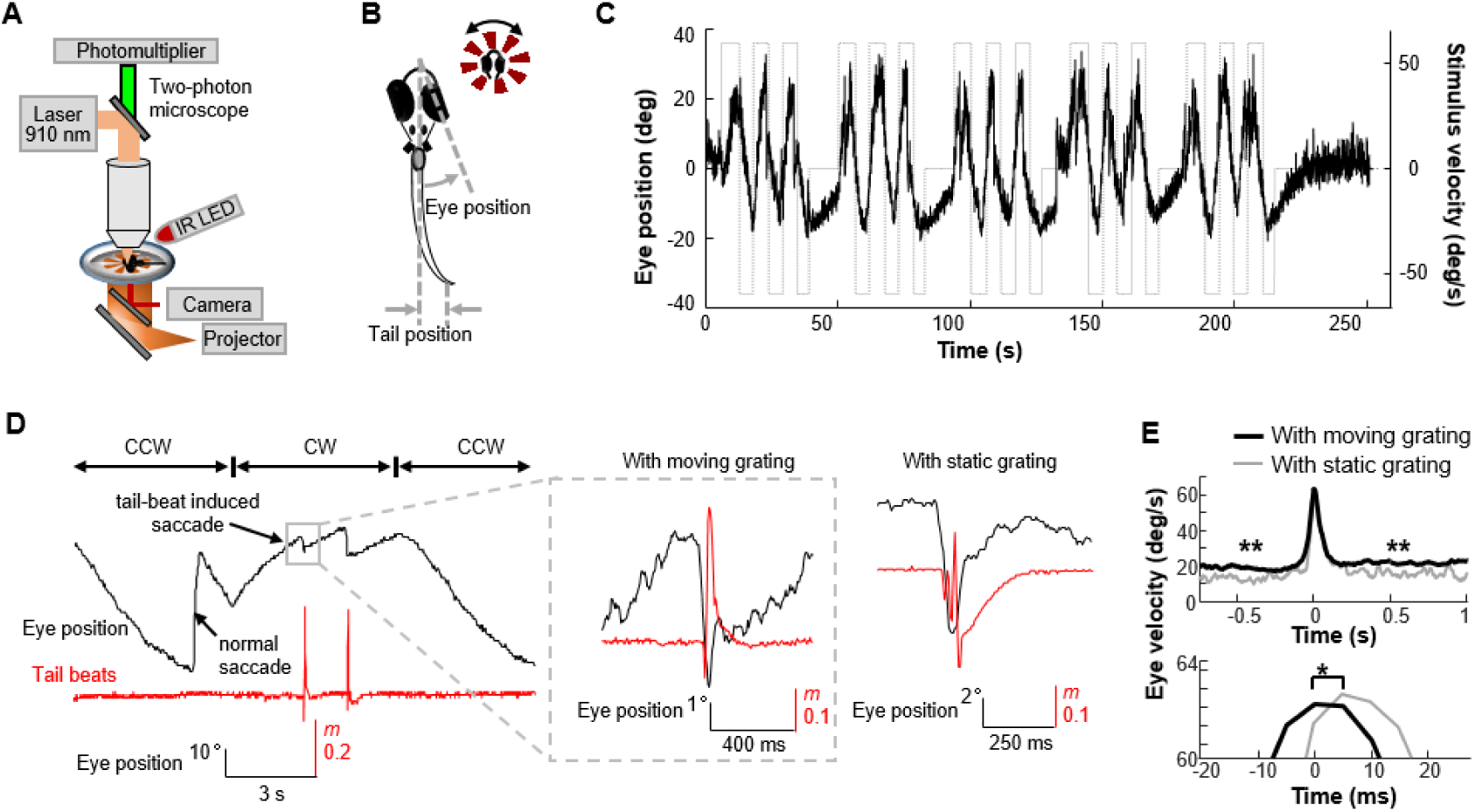
Tail movements modulate optokinetic response (OKR). (**A**) Experimental setup. Zebrafish larvae were restrained in agarose, with eyes and tails (in tail-free condition) free, and placed on a miniature screen which was used for visual stimulation. Video of eye and tail movements was recorded by a fast-speed camera, illuminated by a high-power IR LED near the detective lens of the two photon microscope. (**B**) A radial spinning pattern was presented to the zebrafish larva to induce OKR response. The position of eye was measured as the angle between the long axis of the eye and the midline of the body, while the position of tail was measured as the relative displacement of the tip of the tail. (**C**) The radial grating was rotating with constant velocity and changed direction periodically. Counterclockwise eye positions were defined to be positive. Larval zebrafish tracked the visual movements with a sinusoidal OKR pattern. (**D**) One single beat of tail movement reset the eye position, in opposite to the direction of the ongoing eye velocity, when the visual movement was presented (left panel). The eye continued to move from the new position with same velocity prior to the reset (middle panel). When the visual stimulus was static on the screen, one single beat of tail movement also changed eye position, but eye returned to its previous position soon after the tail movement (right panel). Tail beats were measured as *m(t)*. (**E**) A peak of eye velocity was found associated with a tail-beat. The eye velocities before and after the tail beat were significantly larger when the visual stimulus was moving (upper panel). Latency of the peak was significantly shorter for moving stimulus in comparison with the static ones (lower panel). * P < 0.01, ** P < 0.001.

We found that the slow phase pursuit of the OKR in the larvae was synchronized with the change of direction with occasional fast reset saccades (Figure 1C). In spite of the common OKR patterns, we also found that there were cases where a tail-beat induced a fast reset saccade that was opposite of the ongoing pursuit during the presentation of rotating gratings, and the eye continued moving in the previous smooth pursuit direction after the saccade (Figure 1D, left and middle panels, see SMovie1 for examples). However, the tail-beat induced saccade (TBIS) showed a different pattern when it was evoked during the static grating: the eye returned to its original position by another saccade or by slow drifts (Figure 1D, right panel, see Figure S1_1 for more examples). To evaluate this tail-eye interaction quantitatively, we measured the change of eye velocity around the tail-beat in both rotating and static conditions. The averaged eye velocity showed a significant peak aligned with the onset of the tail-beat, while the baseline velocities before and after the peak were higher in the moving grating condition than in the static grating condition (21.5 ± 0.8 vs. 16.4 ± 1.9 degree/s, mean ± SEM, P < 0.001, before saccade; 24.4 ± 0.7 vs. 20.2 ± 2.7 degree/s, P < 0.001, after saccade. Figure 1E, upper panel). This is consistent with the observation that TBIS during OKR resets the eye position even though the smooth pursuit is resumed after the saccade. Although the peaks of the TBIS showed no difference in amplitude for the two conditions, a significant shorter latency was found for the moving grating condition (2.2 vs. 8.35 ms, P < 0.05, Figure 1E, lower panel and histogram of the latencies: Figure S1_2, P < 0.05, KS test).

This tail-OKR interaction not only reset eye position by the single tail-beat, but also altered the slope of the smooth pursuit when several tail-beats were generated in sequence as a bundle (Figure 2A, see SMovie2 for example). Though the generation of multiple tail-beats varied across individuals in the above-mentioned tail-free condition, the averaged eye velocity of smoot pursuit was significantly smaller than that of the same fish during a tail-immobilized condition (P < 0.05, n = 19, Figure 2B). It is important to note that the head/body of the fish was constrained by agarose and kept stable in both conditions, resulting in a constant visual input signal to the eyes. The lack of tail-induced blurring on visual inputs leaves no space for a feedback control on eye movement from the visual brain areas. The TBIS and its modulation of smooth pursuit during OKR suggested the existence of an efference copy signal of the tail movement upon eye movement control, even when the eye movement was driven by visual stimulus.

**Figure 2.**
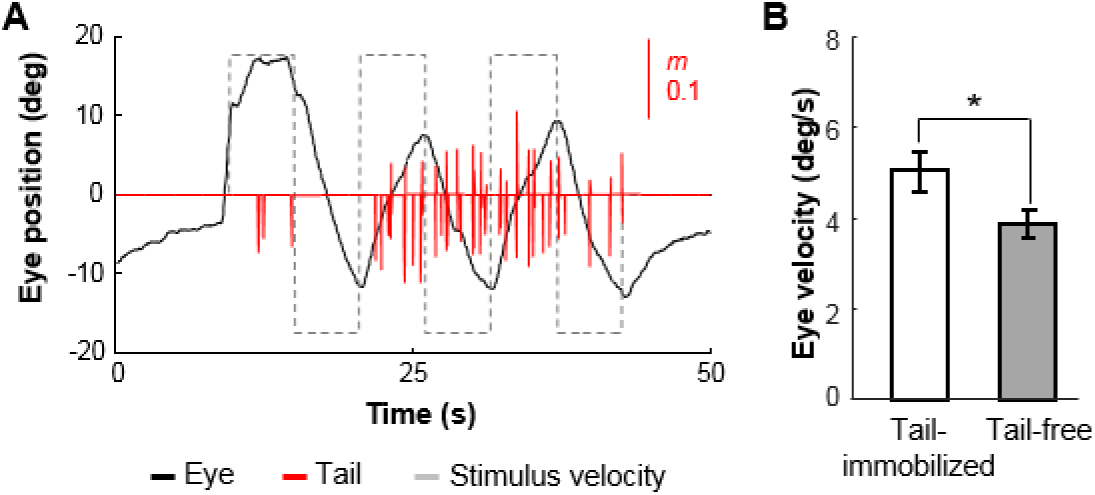
Larger OKR found when the tail is immobilized. (**A**) Occurrence of multiple tail-beats interrupted rhythmic OKR pattern. (**B**) Comparison between eye velocities during the tail-immobilized and the tail-free conditions. For the tail-immobilized condition, the agarose is in contact with tail. The average eye velocity was reduced during the tail-free condition. Error bars indicate SEM; n = 19 fish. * P < 0.05.

### Neural activity in the hindbrain during tail-OKR interactions

To explore the neural basis of the efference copy, including the neurons facilitating the TBIS and its effect on OKR, in vivo two-photon calcium imaging was performed in the tail-free and tail-immobilized conditions. Several studies demonstrated that the neural mechanisms involved in OKR (Portugues et al., 2014), especially the velocity-to-position neural integrator (VPNI) circuit (Miri et al., 2011) and the mechanism for saccade generation (Schoonheim et al., 2010), were located in hindbrain of zebrafish. In this study, we acquired calcium images from hindbrain of zebrafish larvae (*elavl3:GCaMP5G* × *mitfa*^*-/-*^) by a two-photon microscope. For each fish, functional calcium images from one optical section of hindbrain were first acquired for the tail-immobilized condition and then agarose embedding the tail was carefully removed for the tail-free condition, during which the visual stimulus was presented and the behavioral responses were recorded (Figure 1A). Eye positions were determined from the infrared video (Figure 1B) and convolved with an exponential kernel to generate the individualized OKR regressors (Kubo et al., 2014; Portugues et al., 2014). Functional activities were evaluated by pair-wise correlation between calcium traces and OKR regressor, resulting in correlation maps for the two conditions. OKR-sensitive functional clusters were determined by a combination of an automated algorithm (Ahrens et al., 2012) and the correlation maps. The clusters may also be referred as ‘cells’ in general (Kubo et al., 2014) and the calcium responses of each cluster were extracted (Figure 3A). We found a lateralized pattern in the hindbrain where neurons on both sides of the midline responded in opposite phase to OKR (Figure 3A, 3B). When these OKR-sensitive neurons/clusters were pooled across individual fish, the clusters could be grouped into three regions of interest (ROI) based on their spatial coordinates: ROI1 in rostral hindbrain, ROI2 in central hindbrain, and ROI3 in caudal hindbrain (Figure 3C, 3D). Consistent with previous findings, the neurons in ROI1 responded in reversed pace with ROI2 and ROI3 (Figure S3_1, see SMovie3 for example) in a stereotyped manner (Portugues et al., 2014). However, the responses were subjected to change when tail-free and tail-immobilized conditions were taken into consideration, as predicted. More OKR-sensitive neurons were seen in ROI1 and ROI2 for the tail-free conditions, while more neurons in ROI3 were activated in the tail-immobilized condition, demonstrated by density (spatial overlapping) of the neurons (Figure 3C, 3D) or the spatial distribution of the neurons (Figure S3_1B). In spite of the difference in the number of cells (Figure 3E) in the two conditions, the amplitude of the calcium responses measured as averaged correlation coefficients, displayed a similar pattern (Figure S3_1A) with ROI1 and ROI2 are more involved in the tail-free condition, while there is a larger contribution from ROI3 in the tail-immobilized condition (Figure 3F, P < 0.001). It is important to note that the differences in the neural activation in the three ROIs described above in the two conditions were not the direct consequences of tail movements in the tail-free condition. In contrast, brain regions lateral to ROI3 were found to be directly involved with tail beats when a tail regressor was applied to the calcium imaging stacks during the tail-free condition (Figure S3_2).

**Figure 3.**
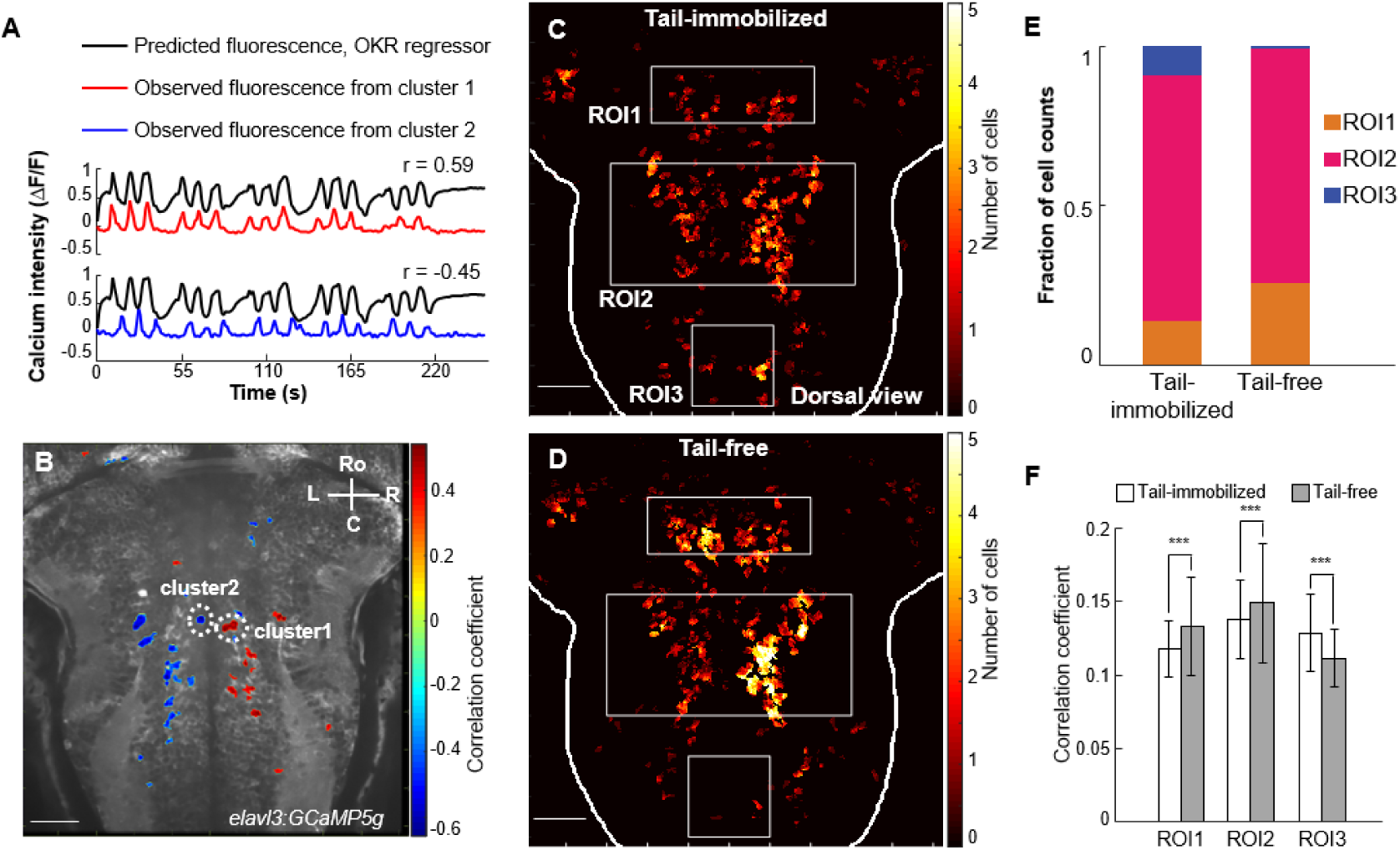
Calcium imaging revealed the involvement of hindbrain during tail-OKR interaction. **(A)** Eye positions were convolved with an exponential kernel using the decay time constant of *elavl3: GCaMP5g* to predict fluorescence (ΔF/F) related with OKR. Example fluorescence traces from two clusters (from **B**, red and blue) showed positive and negative correlations with the regressor, respectively. (**B**) Example of 2D map of image pixels that are correlated with the OKR regressor, from one fish, superimposed on its anatomical reference by averaging images across scans. Note the two clusters (or cells) in white circles have opposite response polarities, as shown in **A**. Ro: rostral; C: caudal; R: right; L: left. (**C**) OKR related neural responses in the tail immobilized condition were pooled together across fish. Pseudocolor scale depicts the number of cells at a given location in the hindbrain of which was significantly associated with OKR. Three regions of interest (ROIs, white boxes) were found: ROI1, rostral hindbrain; ROI2, central hindbrain; ROI3, dorsal-caudal hindbrain. (**D**) Similar response pattern was observed in the tail free condition, with stronger responses in ROI1 and ROI2, and less responses in ROI3. (**E**) Fraction of cell counts of the three ROIs revealed that in the process of tail-OKR interactions, more cells were activated during tail immobilization in ROI3, whereas more cells were activated during tail free in ROI1. (**F**) Correlation coefficient was also different for the two tail conditions in the three ROIs. Larger correlation values were observed in ROI1 and ROI2 during the tail-free condition; larger correlation values were observed in ROI3 during the tail-immobilized condition. Error bars indicate standard deviation. *** P < 0.001.

### 3D function imaging using light-sheet microscopy

To explore the involvement of the entire hindbrain during the tail-OKR interactions and to extend beyond the single slice limitation of two-photon microscopy, a light-sheet microscope was customized for this study. The setup was designed to record 3D calcium signals from zebrafish larvae at a temporal frequency of 1 Hz (Figure S4_1A). A plastic opaque shutter was inserted in the agarose near the eye (Figure S4_1B) to ensure reliable OKR responses elicited by rotating gratings for most individual runs (Figure S4_2). Stacks of calcium images covered most part of hindbrain by 24 images per stack (see SMovie 4 and 5 for demonstration). Datasets from the two tail conditions were co-registered after a volume-based correction for motion artifacts and normalized to the Z-Brain Atlas template brain (Randlett et al., 2015) by an affine transformation. Three regressors were generated in the same manner as in the two-photon experiments: the OKR regressor from the eye positions, a stimulus position regressor, and a saccade regressor (Figure S6). Functional activation maps were calculated by measuring the maximum correlation coefficients between the calcium responses and the regressors (Figure 4B, also see SMovie6 for 3D example). Group level analysis on the functional maps revealed brain regions involved in OKR were similar to that found in the two-photon experiments, including the rostral hindbrain (rHB), the central hindbrain and the dorsal-caudal hindbrain (dcHB, Figure 4C). It is apparent that the activations in the tail-immobilized condition are stronger in dorsal hindbrain and extend to further caudal regions. Contrast analysis of the two conditions by a paired t-test at group level confirmed this observation by demonstrating that dcHB has stronger activations in the tail-immobilized condition while the deep rHB is more involved in the tail-free condition (Figure 4D). The results coherently reproduced the pattern found in the two-photon experiments, even though the details of the visual stimuli and the imaging setup were different in many aspects. The capacity of volumetric imaging provided by the light-sheet microscopy not only facilitated the normalization of each dataset to the ZBrain Atlas (Randlett et al., 2015), hence helping artefact correction for individual runs and for group level tests, but also enabled precise localization of brain activations to well-established anatomical brain structures (Figure S4_3). The tail-free related rHB clusters were recognized as within the anterior cluster of nV trigeminal motorneurons, Vglut2 Cluster 1, and Gad1b Cluster 1. Meanwhile, the dcHB clusters for the tail-immobilized condition were found to be scattered among Gad1b Stripe 2, Vglut2 Stripe 3, and noradrendergic neurons of the interfascicular and Vagal areas (Figure S4_4). The rHB clusters have been demonstrated to be related with saccade and tail movements during OKR (Portugues et al., 2014). The dcHB clusters are within the hVPNI areas (Miri et al., 2011). In combination with behavioral results, two-photon experiments and light-sheet calcium imaging data suggested that the enhanced rostral activations and suppressed dorsal-caudal activations for the tail-free condition may originate from a push-pull signal from the tail movement center to the saccade generating circuit and the VPNI circuit. It is important to note that in spite of the double dissociation pattern observed in the two brain regions (F(1, 40) = 29.3, P < 0.001), the averaged coefficient in rHB clusters is significantly smaller than that in dcHB clusters for the tail-immobilized condition (P < 0.001) while for tail-free condition the two brain regions showed almost the same level of responses (Figure 4E). The results fit with the proposition that the responses of the saccade generating circuit in the rostral hindbrain only control the fast-phase (saccade) of the OKR (Schoonheim et al., 2010), thus show smaller correlations to the eye position traces, while the VPNI circuit in the dorsal-caudal hindbrain determines the slow-phase of the OKR and has larger correlations to the eye position in general (Miri et al., 2011) for the tail-immobilized condition. In the tail-free condition, the fact that tail-beats induced extra saccades implied that the tail movement signal changed the neural activity in hindbrain in presence of evidence: 1) it increased the responses of saccade generating neurons in the rHB, and 2) it inhibited the VPNI mechanism in the dcHB.

**Figure 4.**
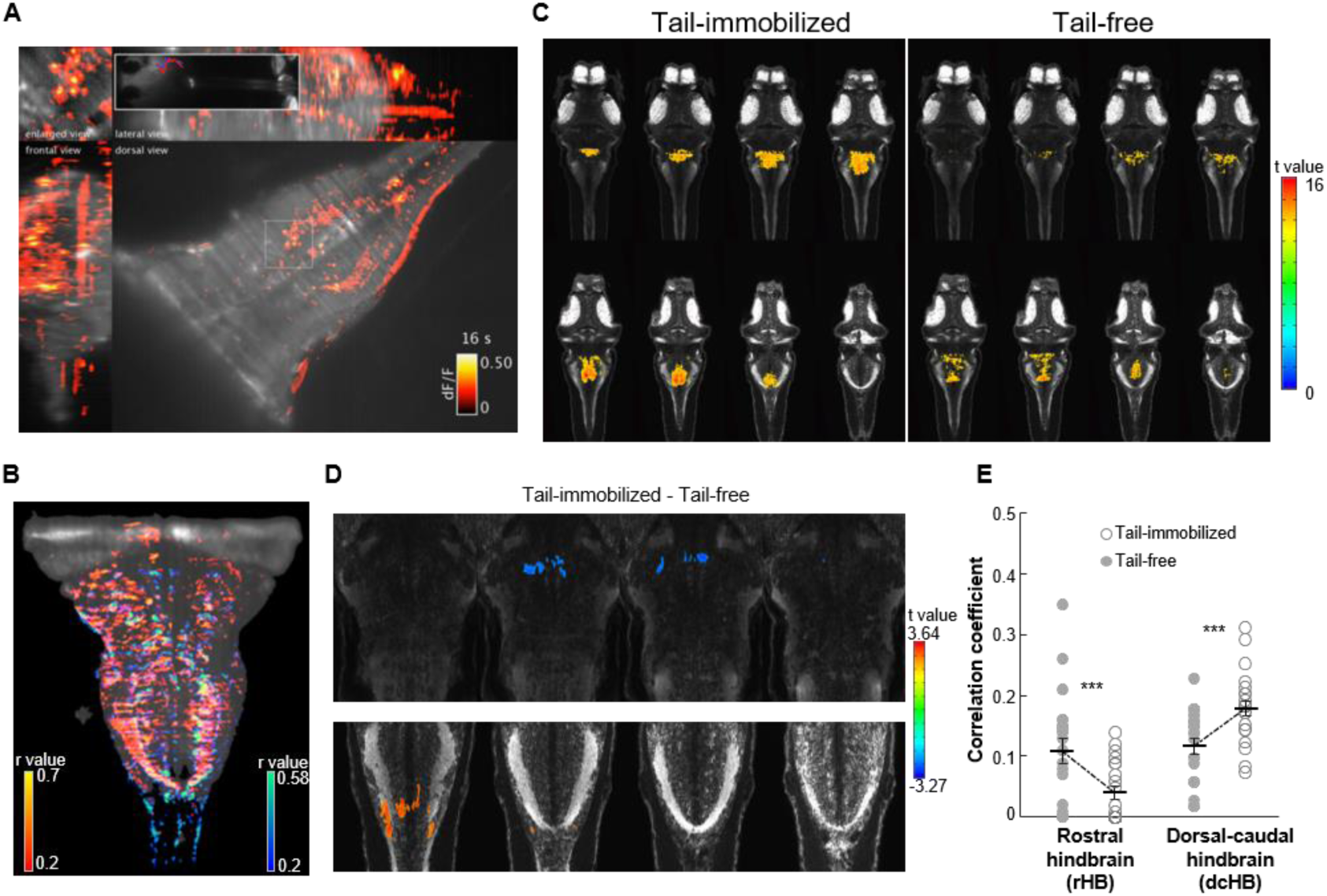
Calcium imaging by light-sheet microscope provides a volumetric map of the neural activations during tail-OKR interaction. (**A**). Frontal, dorsal and lateral projections of volumetric imaging of calcium activity (ΔF/F) at hindbrain during tail-OKR interaction, acquired by a custom light-sheet microscope. Left top corner, enlarged view of the region outlined by white box in dorsal projections. Inset, infrared videos of the fish, with traces of eye position superimposed. **(B)** An example from one larva, showing active neural populations involved in OKR. Pseudocolor scale, correlation coefficient with OKR; Red, tail-free; blue, tail-immobilized. (**C**) Group averaging, after mapping individual volumetric data to a zebrafish atlas (Z-Brain Atlas), demonstrated similar activity pattern across both tail-free and tail-immobilized conditions. Brain regions including the rostral hindbrain, the central hindbrain and the dorsal-caudal hindbrain. (**D**) Group contrast revealed a stronger response in the rostral hindbrain for the tail-free condition (upper panel) and larger response in the dorsal-caudal hindbrain for the tail-immobilized condition (lower panel). These brain regions, indicating the neural substrates of the tail-OKR interaction, in consistent with the findings by two-photon imaging in Figure 2. (**E**). Degree of involvement (correlation coefficient) showed a double dissociation between the two brain regions (rostral and dorsal-caudal hindbrain) in correspondence to the two tail conditions. This result indicates specific roles of the brain regions regarding tail-OKR interaction. Each dot represents data from one brain region of one fish in a given tail condition. Error bars indicate SEM. n = 22 fish. *** P < 0.001.

### Information flow revealed by Granger causality

The double dissociation pattern of functional activities may reveal an intrinsic push-pull signal on rostral and dorsal-caudal hindbrain respectively, but it could also be explained by larger variability of eye traces during the tail-free condition due to the extra TBIS, while the neural responses in the hindbrain kept the same. To address this question, we utilize an independent approach to explore the alterations of hindbrain neural dynamics under the two conditions. We calculated the information flow, measured as Granger causality (Granger, 1969), between the rHB/dcHB clusters and other parts of the hindbrain. The information flow between two signals/time series has been determined by estimating/forecasting the signals with a univariate autoregressive model or with a multivariate autoregressive model while taking another time series into consideration (Figure 5A). We evaluated the information flow projected from other hindbrain regions into rHB/dcHB clusters (or vice versa) by paired t-test on Granger values at group level separately. The results showed that other hindbrain regions exchanging information with rHB/dcHB are located mainly within the central hindbrain (Figure 5B). However, these regions in the central hindbrain are more lateral to the activations found in the correlation maps in Figure 3C. More importantly, the tail conditions significantly altered information flow in the hindbrain: in cases were information flows into rHB, there were more voxels in the hindbrain involved during the tail-free condition, whereas in the tail-immobilized condition, there were more voxels in the hindbrain receiving information from dcHB. This pattern was consistent across a wide range of thresholds (Figure S5. P < 0.05, into rHB; P < 0.002, from dcHB, KS-test). It confirmed the proposed push-pull mechanism in the hindbrain, since the information flow measured as Granger causality is irrelevant to how we define the functional maps. Anyhow, we also tested the links between averaged Granger values and the coefficients of OKR activation for every rHB/dcHB cluster by Pearson correlation at group level with individual data. We found that only two clusters in the dcHB showed significant positive correlation between the strength of information flow projecting to other parts of the hindbrain and the coefficients of OKR activation (Figure 5C and 5D, r = 0.32, P < 0.05). Fish larvae with higher OKR activation in dcHB projected stronger information to the central hindbrain. Due to the well-defined functional meaning of the OKR regressor, it is reasonable to speculate that the clusters in dcHB, possibly part of the VPNI circuit, play a leading role in the tail-OKR interaction in the hindbrain.

**Figure 5.**
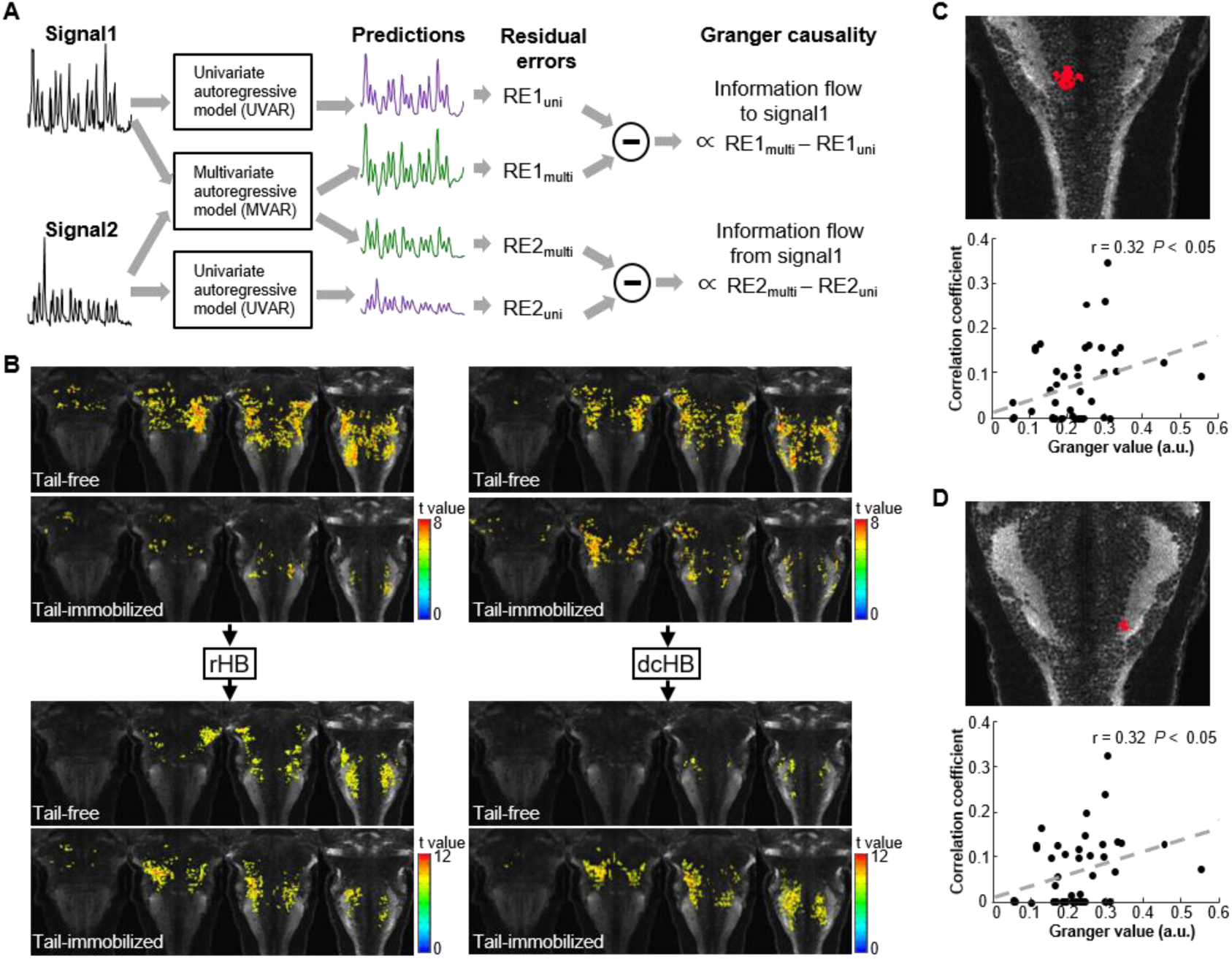
Information flow measured as Granger causality. (**A**). A schematic illustration of the procedure in measuring information flow as Granger causality between two signals/time series. A time series could be estimated by a univariate autoregressive model or, with the existence of another time series, by a multivariate autoregressive model. To what extend the residual errors were reduced in the multivariate model compared with that of the univariate model, is defined as Granger causality, a measurement of the information flow from the helper time series to the signal to be estimated. (**B**) Maps of information flow between rHB/dcHB and other parts of hindbrain for the two tail conditions. In the comparison of the two conditions, there are more brain areas project information to rHB in the tail-free condition and dcHB casts information flow to larger brain areas in the tail-immobilized condition. (**C**) and (**D**) Two clusters (upper panels) in dcHB showed positive relations between their information flow (Granger values) projected to other brain areas and the functional activations (correlation coefficients) of OKR (lower panels, pooled across two tail conditions), indicating a link between the information flow sourcing from dcHB and its functional role during OKR. Each dot represents one fish in one condition.

### Single neuron dynamics and covariance with different regressors confirmed the push-pull mechanism

There are several direct predictions that could be derived from the proposed push-pull mechanism.

Firstly, the activity of the saccade generating circuit before TBIS determines the neural dynamics after it. These neurons, driven by visual stimulus, generate periodic output when the accumulated inputs reach a threshold (Schoonheim et al., 2010). With the help of this background activity, the efferent signals of the tail push the response of these neurons to surpass the threshold faster and earlier, compared with the situation where the stimulus is static and the background activity is missing (Figure 1E). To test this idea, the calcium responses from saccade-related neurons (see Methods for details) were sorted into short clips/episodes for each saccade and averaged into three categories: TBIS with moving grating, TBIS with static grating, and normal saccade without tail-beat. As predicted, the averaged calcium intensity for the TBIS with moving grating is significantly larger than that of the TBIS with static grating, before the onset of the saccade (P < 0.05, Figure 6A). This difference is more than likely due to accumulated information of the moving grating, since the normal saccade without tail-beat also showed a significantly larger signal before the onset of the saccade (P < 0.05). When the episodes were sorted by the calcium intensity at the onset of the saccade, it is also obvious that the larger the responses before the saccade, the larger calcium signal intensity at the onset of the saccade (Figure 6B). The baseline activity before the saccade not only determined the responses at the onset of the saccade, but also influenced the peak latency of the neural dynamics of these neurons, which is revealed by the comparison of the averaged curve of the first half of the episodes with that of the second half of the episodes (Figure 6C). For TBIS with moving grating and normal saccades with moving grating, the larger baseline activity leads to earlier peak of the neural dynamics. However, even though averaged curves for the episodes of the TBIS with static grating were generated by the same procedure, the peak latency is the same for the first half and the second half of the episodes, indicating different neural dynamics without tail-OKR interactions. It is interesting to note that the shorter peak latency for TBIS with moving grating, in comparison with normal saccades with moving grating (Figure 6A red vs. green, Figure 5C blue curves in left and right panels), also confirmed the proposed pull mechanism from tail movement for saccade generating during OKR.

**Figure 6.**
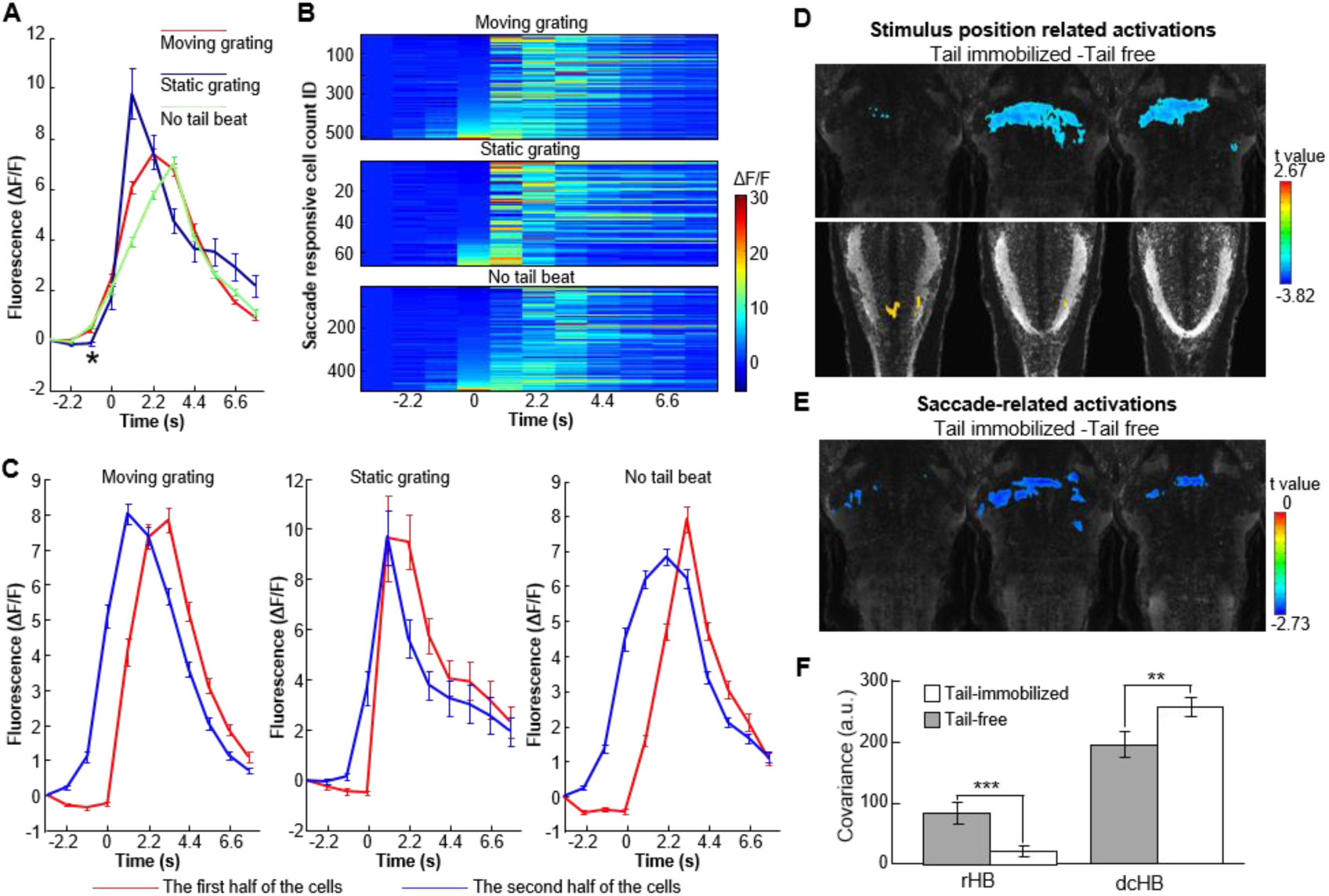
Model predictions evaluated by single neuron dynamics and covariance with different regressors. (**A**). Averaged calcium responses from neurons activated by saccades. The neural responses are significantly higher before the onset of the saccades when the stimulus is moving, disregard of the saccade is tail-beat induced (red) or OKR induced (green), in comparison with the saccades induced by tail-beat when the grating is static (blue). (**B**). Individual activity trace of neurons activated by the three types of saccade respectively, sorted by the activity at the onset of the saccade. Notice the responses before the onset of the saccades for the two types during moving grating (upper and low panels) are negatively correlated with the peak latency of the calcium signal. (**C**). For each of the three types of saccade, the sorted individual activity traces in (**B**) were averaged for the first half and the second half of the traces separately. The second half of the traces showed higher amplitude before saccades. Higher amplitude before saccade onset leads to shorter peak latency when the stimulus is moving (left and right panels), but no such relation was found for the TBIS when the grating is static (middle panel). (**D**). The activations for stimulus position regressor showed larger responses in dorsal-caudal hindbrain and smaller responses in rostral hindbrain for tail-immobilized condition. (**E**). For activations related with saccades, rostral hindbrain was found to be more active in tail-free condition. (**F**). Covariance of the calcium signals defined by stimulus location regressor indicates an enhancement in rostral hindbrain; covariance of the calcium signals defined by saccade regressor indicates an inhibition in dorsal-caudal hindbrain by the tail beats. * P < 0.05, ** P < 0.02, *** P < 0.01.

Secondly, it is predicted that the correlated calcium activity with the stimulus position regressor would be different for tail-free and tail-immobilized conditions in dcHB, if the tail movement inhibits the VPNI circuit thus inhibit the information integration of the visual inputs. That is exactly what we found in our light-sheet datasets (Figure 6D). As expected, in the tail-immobilized condition, calcium activity showed higher correlation with the stimulus position regressor than in the tail-free condition. Since the stimulus position regressor is the same for both conditions (Figure S6), the only explanation is that the tail movement inhibited the neural activity in the tail-free condition, as a push mechanism. The covariance of the calcium signals in dcHB was also consistent with this prediction that smaller covariance was found for the tail-free condition (Figure 6F).

The third prediction is that although the push-pull signal projected to rHB and dcHB from the same source, possibly the CPG center for tail movement, it required a local circuit to generate the extra resetting saccade (Schoonheim et al., 2010). Thus, for the tail-free condition, the neural activity in rHB clusters had more saccade-related components than that in the tail-immobilized condition, but in dcHB no such difference is necessary. The correlation maps with saccade regressor demonstrated this prediction (Figure 6E). The covariance was also significantly larger for the tail-free condition in rHB (Figure 6F), while there is no difference found in dcHB.

## DISCUSSION

We have demonstrated that tail-beats could induce extra saccades when larval zebrafish were presented with rotating gratings. This suggests an interaction between tail-beat and OKR, most likely due to an efference copy signal from the tail-movement center to help stabilize visual perception. Calcium imaging via both two-photon microscopy and light-sheet microscopy revealed that the rostral hindbrain was more active during the tail-free condition while the dorsal-caudal hindbrain responded stronger during the tail-immobilized condition. The different neural responses for the two conditions suggested a push-pull mechanism for the tail-OKR interaction in the hindbrain.

### Efference copy resets eye position during OKR

A general framework to understand motion perception and motion control is the perspective of internal model (Lisberger, 2009). The principal concept is that sensory information is an afferent signal transferred from peripheral sensors to central processing units. The central nervous system holds a mechanical model of the motion objects/the environment, whose dynamics generates proper motor commands and helps to predict future events. To maintain a stable representation of the environment, the neural system needs to cope with the self-generated noise/artefacts (reafference signal) generated by its own movements. It is supposed that the motion center not only generates motor commands to the motor system, but also sends duplicated ones, termed as efference copy by von Holst or corollary discharges by Sperry (Lisberger, 2009; Sommer and Wurtz, 2002, 2008), to the sensory system for predicting the forthcoming changes. This prediction is compared with the reafference signal to keep a stabilized perception and maintain a sustained motion control (Shadmehr et al., 2010), as well as increases the signal-to-noise ratio of the sensory system (Frens and Donchin, 2009; Lisberger, 2009; Sommer and Wurtz, 2008). The existence of efference copy was first demonstrated by the suppressed sensory signals located at the level of afferent fibers and/or the central neurons, in the mechanosensory system of the crayfish (Edwards et al., 1999; Kennedy et al., 1974) and electrosensory system of the electric fish (Bell, 1981). Further evidence from the vestibulo-ocular reflex (VOR) in non-human primates suggested that during active or passive vestibular head movements, the activity of vestibular nucleus was suppressed (Roy and Cullen, 2001) when the motor-generated expectation matches the activation of proprioceptors in the neck (Roy and Cullen, 2004). Although it is hypothesized that the efference copy for this kind of VOR estimation arises from the vestibular system (Lisberger, 2009), the effect could also be explained by coordinated timing of motor commands (Braitenberg et al., 1997; Llinas, 1988). The latter idea was supported by the fact that the delay in eye-head coordination is nearly 0 during passive whole-body or self-generated head movements in the guinea pig (Shanidze et al., 2010a; Shanidze et al., 2010b). In addition, several other animal species also shows synchronized body/head-eye movements while studying the visual perturbation during locomotion. This indicates a direct contribution to eye movement control by head/body motor commands (Chagnaud et al., 2012). Using a variety of in vitro and in vivo preparations of Xenopus tadpoles, Lambert et al. demonstrated that this conjugate eye movements, in opposite to horizontal head displacements during undulatory tail-based locomotion, was produced by the spinal locomotor CPG derived efference copy (Lambert et al., 2012).

In the current study, we found that larval zebrafish with head and body embedded in agarose could generated extra saccades that was induced by tail-beats during their perception of whole-field rotating visual stimulus. This is the first direct evidence showing that efference copy could drive compensatory eye movements during active visual perception. During a single tail-beat, the induced extra saccade resets the eye position to the opposite of the OKR direction, and the latency is even shorter than that of the TBIS when the visual stimulus is static. Moreover, when multiple tail-beats were generated in a sequence, there was a reduction in OKR amplitude. These facts would have been overlooked if solely explained by synchronized motor commands or timing coordination, suggested a more functional relevance of the tail-related efference copy in visual perception and visual stabilization.

### The rostral hindbrain combines visual information and tail signals for saccade command

The first observation of our calcium imaging, consistent across the two-photon imaging experiment (Figure 3) and the light-sheet functional results (Figure 4), is that neurons in rHB showed a stronger response in the tail-free condition than in the tail-immobilized condition. These neurons are within rostral hindbrain areas that are related with eye and tail movements (Portugues et al., 2014). Since the spatial distribution of these neurons are different from the active neurons that are directly linked with tail movements (Figure S3_2), these rostral hindbrain neurons are mostly the neural underpins of the tail-OKR interaction, but not a direct consequence of the tail movements. This is also confirmed by the information analysis showing that the rHB received information from a broader area of the hindbrain in the tail-free than in the tail-immobilized condition (Figure 5), indicating a role of information integration in the rHB. Though proposed as a tool for economic data analysis (Granger, 1969), the Granger causality used here has been successfully applied in human functional brain research (Roebroeck et al., 2005), neurophysiology of primate visual perception (Gregoriou et al., 2009), zebrafish functional analysis at neuron level (Fallani Fde et al., 2015) and system level (Rosch et al., 2018). In this study, Granger causality revealed that the rHB is a tail-OKR interaction center when the tail is free to move during visual driven eye movement. When a certain threshold has been reached, accumulating activations in these saccade preparation areas (Wolf et al., 2017) would trigger saccade commands which are projected to saccade generators (Schoonheim et al., 2010) and oculomotor integrators (Goncalves et al., 2014). A direct prediction of this assumption is that this threshold would be reached easier when the tail is free during the viewing of a moving stimulus, resulting in shorter latency to peak responses after the saccade. We found the exact pattern in the single neuron dynamics in the two-photon experiments. For saccades present during the moving stimulus, non-dependent on tail-beat-induction, had larger neural activities before the onset of the saccades than the TBIS without a moving stimulus (Figure 6A). It demonstrated the preparatory neural activity in the rHB that is related to moving visual inputs, which could be recognized as OKR-related components (Portugues et al., 2014). Moreover, the trial-by-trial neural dynamics revealed a shorter latency of the peaks for the TBIS over normal saccades, indicating an integration of tail signals into the on-going visual inputs (Figure 6C).

It is interesting to note that when neural activities measured by saccade regressors were compared, the rHB also showed enhanced correlations with the saccade regressor in the tail-free condition compared to the tail-immobilized condition (Figure 6E). This difference, from another perspective, evidently demonstrates that the neural activities in the rHB clusters are not the final step to determine the behavioral detectable eye movements, otherwise the correlations between neural responses and the saccade regressors would be equal in both conditions.

### Suppressed VPNI circuit during tail-OKR interaction

We found suppressed activity in dcHB in the tail-free condition. It is within the hVPNI brain regions (Daie et al., 2015; Miri et al., 2011; Portugues et al., 2014). We believed that this difference is due to the inhibition of the efference copy from the tail motor center in the tail-free condition. It is consistent with the inhibitory role of efference copy to compensate for the reafferent sensory input and to help detect changes in the environment during self-generated movement (Lisberger, 2009; Sommer and Wurtz, 2002, 2008), under the topic of VOR (Lisberger, 2009; Roy and Cullen, 2001, 2004) and other movements (Shadmehr et al., 2010). Moreover, there are also evidence that higher level of perception, such as space (Ross et al., 1997) and time (Winter et al., 2008) are transiently distorted around the moment of a movement. In the current study, dcHB not only showed smaller correlation with the OKR response in the tail-free condition (Figure 3 and 4), but also showed smaller correlation with stimulus position regressors, possibly due to inhibited VPNI circuits (Figure 6D). The VPNI integrates inputs from upstream visual and vestibular information and serves as a suitable plant for internal model of motion integration. However, we can’t rule out the possibility that the sensory suppression may be achieved within more peripheral/lower-level neural circuits, such as pretectum (Kubo et al., 2014), but suppressing VPNI circuit activity is definitely a more effective approach, if the transient change of visual and proprioceptive inputs induced by the extra resetting saccades are taken into consideration.

### A third approach for efference copy to interact with ongoing motion perception

Previous studies have demonstrated that there are at least two approaches for efference copy to modulate internal model: the efference copy interacts with direct representation of sensory information, either by sensory suppression (Lisberger, 2009) or remapping (Wurtz, 2018); the efference copy coordinates compensatory motor patterns to eliminate sensory reafference (Chagnaud et al., 2012). Here we suggest a third approach that the efference copy may regulate the neural activities of secondary cognitive modules, a kind of state estimators (Frens and Donchin, 2009) for motor preparation and sensory information integration during active motion perception, in a push-pull manner (Figure 7). As demonstrated in zebrafish larvae during tail-OKR interaction, the visual motion signals are projected from pretectum to dcHB. In dcHB, the VPNI mechanisms generate the necessary information for predictive eye positions. In central hindbrain (cHB, including ABN), the VPNI signal and the saccade command from rHB triggered the eye movements. In the meantime, there are also projections from dcHB to the central pattern generator (CPG) in spinal cord. When the tail is immobilized in the agarose, the CPG ceased to generate a motor command, most likely due to the mismatch of predictive sensory feedback from the tail (Grillner et al., 1998; Roy and Cullen, 2004), thus no efferent signal is sent from CPG to cHB and dcHB (Figure 7A). When the tail is free to move, the CPG generates motor commands for tail beats during the OKR response, and also sends efferent signals to cHB and dcHB. The excitatory signals from CPG to cHB are summed with the velocity information from the visual inputs (Wolf et al., 2017) and contribute to a higher information flow from cHB to rHB to ramp up the saccade commands. Meanwhile, the efferent signals from CPG to dcHB, especially the VPNI neurons, are inhibitory and reduce the information flow from dcHB to cHB (Figure 7B). Since the sensory system and motor system are occupied by ongoing motion processing, it is a reasonable better choice for efference copy to modulate on these secondary integrative modules (state estimators) as a third approach.

**Figure 7.**
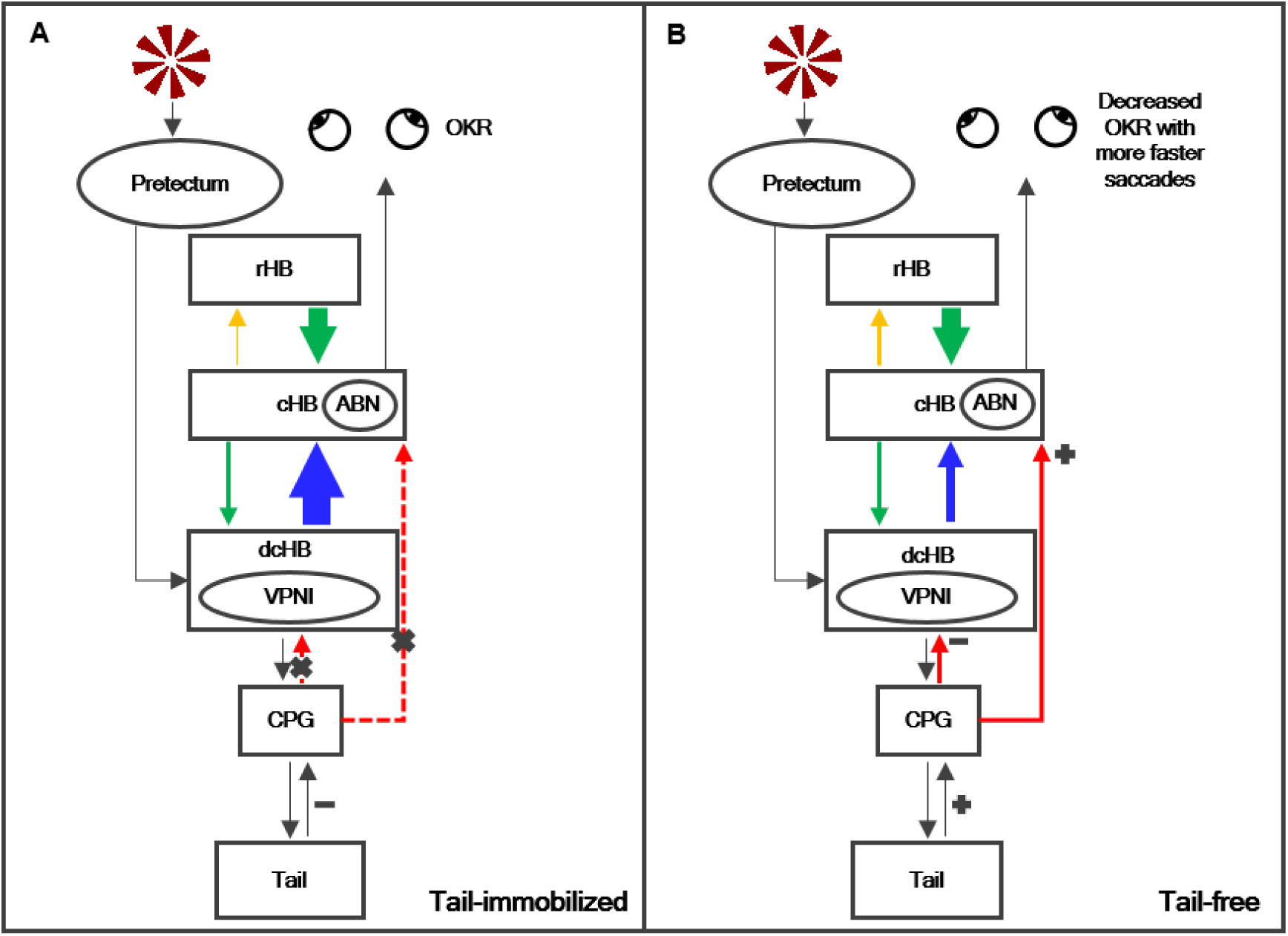
Schematic model for the third approach of efference copy functioning. (**A**) In tail-immobilized condition, there is no efference copy involved. (**B**) When the tail is free to move, the CPG project efference copy to hindbrain via a push-pull manner, on saccade preparation module and VPNI circuits. Orange, enhanced Granger causality in the tail-free condition; blue, inhibited Granger causality in the tail-free condition; green, stable information flow regardless of tail conditions. The width of the lines are proportional to size of the brain regions involved. Red, efference copy from CPG to hindbrain.

### Zebrafish as a good candidate for internal model research

The key point of the internal model is that the neural representations of motion events around the animal and its motor controls in response to the changing environments have intrinsic dynamics, probably due to the manifold constraint of the neurons (Sadtler et al., 2014). The system dynamics follows the same kinetic of the real world, predicting the coming events and correcting behavioral errors related with evoked/self-generated movements (Berkes et al., 2011). This capability, probably inherited from evolutionary adaptation as a neural resonance in response to the real world (Gibson, 1972), not only helps the animal to cope with changes in the environment in a pre-defined manner (Green and Angelaki, 2010; Lisberger, 2009), but also believed to enrich higher level perception such as internal monitoring (Shadmehr et al., 2010), or mirror neural system (Kilner et al., 2007). The relevant studies are mainly based on mammals, but are now expanded to vertebrate zebrafish. The advantage of zebrafish is the translucent brain that enables optical imaging (Ahrens et al., 2012) and optogenetic manipulations (Arrenberg et al., 2009; Goncalves et al., 2014) with the help of genetic tools (Neuhauss, 2003; Renninger et al., 2011). Several neural circuits related with internal model have been explored, e.g., motor adaptation (Ahrens et al., 2012), threat assessment and prey detection (Barker and Baier, 2015; Bhattacharyya et al., 2017; Del Bene et al., 2010; Dunn et al., 2016; Semmelhack et al., 2014; Temizer et al., 2015), behavioral context of short-term memory (Daie et al., 2015), sensory motor integration (Knogler et al., 2017; Koyama et al., 2011; Mu et al., 2012; Schoonheim et al., 2010; Wolf et al., 2017; Yao et al., 2016), OKR (Kubo et al., 2014; Portugues et al., 2014), VPNI (Goncalves et al., 2014; Miri et al., 2011), motion after effect (Perez-Schuster et al., 2016), and internal rhythm (Kaneko et al., 2006; Romano et al., 2015; Sumbre et al., 2008; Warp et al., 2012; Wyart et al., 2009). With the advancement of optical imaging methods, the current study contributes a small yet important piece of the neural representation of the internal model: the role of efference copy on the tail-OKR interaction and a push-pull mechanisms in hindbrain to support it.

More than that, there is evidence that the disorder of body movement system leads to the constraining eye movement in patients with Parkinson’s disease (Ambati et al., 2016), which implies a common or interactive system to control eye and body movement simultaneously (Srivastava et al., 2018). The current study may also shed lights on the potential clinical anchor points of the disorders in locomotion-eye coordination with zebrafish model (Huang and Neuhauss, 2008).

## Acknowledgments

We are grateful to Dr. Drew Robson and Dr. Florian Engert for providing *elavl3:GCaMP5g* line, Dr. Jiulin Du for providing the *Nacre* (*mitfa-/-*) line, China Zebrafish Resource Center (CZRC) for providing the *AB* wild type line. We also thank Ms. Yan Teng for two-photon imaging technical support, Ms. Kun Hu and Xin Zhou for behavioral experiment preparations.

This work was supported in part by the Ministry of Science and Technology of China grant (2015CB351701, 2012CB944504), the National Nature Science Foundation of China grant (31730039, 91132302),and the Chinese Academy of Sciences grants (ZDYZ2015-2, XDBS32000000, XDB02010001, XDB02050001, KSZD-EW-Z-001).

## Author contributions

SS, WZ and LZ conceived the experiments. LC, WZ, KRD and LZ supervised the study. SS, and LZ performed behavioral and two-photon imaging experiments. SS and CQ performed light-sheet imaging experiments. SS, ZZ, MMM and LZ analyzed the data. SS, ZZ, MMM, WZ, KRD and LZ prepared the manuscript.

## Data availability

All data and codes used for the analysis are available from the authors on request.

## METHODS

### Animals

Adult zebrafish (*Danio rerio*) are maintained at 28°C under 14/10 day/night cycle. All embryos and larvae are raised in the E3 embryo medium (60× E3B: 17.2g NaCl, 0.76g KCl, 2.9g CaCl_2_.2H_2_O, 4.9g MgSO_4_.7H_2_O dissolved in 1 L Milli-Q water; diluted to 1× in 9 L Milli-Q water plus 100 μl 0.02% methylene blue) (Sumbre et al., 2008). Larvae used in this study are offspring of *elavl3:GCaMP5G* transgenic fish and the *mitfa*^*-/-*^*(nacre)* mutant fish, age between 5 - 7 days post-fertilization (dpf).

### Larvae Preparation

Zebrafish larvae were embedded dorsally in 1.8% low-melting-temperature agarose made with embryo medium at the center of a glass-bottom cell culture dish (Nest801002, outer diameter 35 mm, inner diameter with cover glass 15 mm, Figure 1A). The agarose was stored at 44°C before applying to the culture dish (Bianco and Engert, 2015). The bottom surface of the dish was covered by light-diffusing screen film. A rectangular window, which is slightly larger than the size of fish larvae was opened at the center of the dish, allowing the image of the fish being captured from below by an infrared camera, while the screen film itself served as a projector screen for visual stimulation. The agarose around the eyes and tail (in tail-free conditions) was removed allowing free movement. The same setup was utilized for both behavioral tests and two-photon imaging experiments.

In light-sheet calcium imaging experiments, zebrafish larvae were embedded onto a plastic stage (Figure S4_1A). The top surface of the stage was covered by a piece of light-diffusing screen film with a proper rectangular window. The agarose was removed from around the eyes and tail (in the tail-free condition). The transparent plastic stage was placed in a PMMA (acrylic glass) specimen holder/chamber filled with embryo medium and was perpendicular to the illumination path (Figure S4_1B). The piece of the specimen chamber facing the illumination objective lens was replaced by a glass cover slip. The whole chamber was positioned on a 4-axis (xyz + pitch) manual positioning stage. All animal experimental procedures followed the guidelines of the Institutional Animal Care and Use Committee at the Institute of Biophysics of the Chinese Academy of Sciences (Beijing, China).

### Visual stimulus

To elicit the OKR (optokinetic response) eye movements, a rotating grating of fixed angular velocity was used, which consisted of radial dark and light stripes with angular velocity of 60 degrees/second and 1/45 cycle per degree spatial frequency. Each run of the visual stimuli contains 5 sessions. Each session included 3 clockwise and counter-clockwise cycles of rotations for 33 seconds, followed by a static grating for 11 seconds. During each cycle, the grating rotates clockwise for 5.5 seconds and counter-clockwise for another 5.5 seconds (Figure 1C). The visual stimuli were generated by Matlab (Matlab 2011a, MathWork) and Psychtoolbox (PTB-3) presented by a projector (GP1, BenQ Corporation, China) with its lens system customized for short focus distance and small field-of-view. In addition, only the red light was enabled on the projector. The same setup was used for both behavioral experiments and two-photon imaging. For behavioral experiments, each fish was tested with 10 runs, with freely moving tail. For two-photon imaging experiment, the fish was tested for 2 runs with agarose around the tail (tail-immobilized condition), and was scanned for another 2 runs after the agarose around the tail was carefully removed (tail-free condition).

During the light-sheet imaging experiment, the sequence of stimulus was different from above: each run of the stimulus had 5 clockwise and counter-clockwise cycles of rotations for 55 seconds while the static grating remained for 11 seconds. The fish was scanned for a single run in tail-free condition, and agarose was added to immobilize tail before another run as tail-immobilized condition. Visual stimuli were presented by a projector (Model X2, Coolux, Shenzhen, China) with customized lens system and light source modifications as mentioned above (Figure S4_1A).

### Light-sheet optical setup

The light-sheet was obtained by rapidly scanning a focused laser beam (Figure S4_1A). A 488 nm 20 mW Coherent Sapphire laser beam was projected onto a two-axis galvanometric scanning mirror (Century Sunny TSH8203MAC). The x-axis was driven sinusoidally at 200 Hz to create the light-sheet. The z-axis was programmed to stop at 24 possible angles in turn during 1 s to enable vertical displacement of the light-sheet. The angular deflection of the incident light was transformed into a horizontal/vertical displacement by a scanning lens (Thorlabs CLS-SL) then refocused by a tube lens (Thorlabs ITL200) onto the entrance pupil of a long working distance semi apochromat objective (Olympus XFLUOR4X/340). The thickness profile of the light sheet was 8.3 um, measured as previously described (Panier et al., 2013).

The detection arm was equipped with a high-NA (0.8) 16x water-immersion long working distance objective (Nikon LWD 16x WD 3.0) mounted onto a piezo nanopositioner (PI P-725). Fluorescence was collected by a tube lens (Thorlabs ITL200) and passed through a notch filter (Thorlabs NF488-15) and a custom low pass filter (550 nm), before the image was captured by a CMOS camera with the resolution of 2560 × 2160 pixels. The two filters eliminated 488 nm photons, the red light of the visual stimulus projector and the infrared illumination used for the behavioral recording. A custom-made software acquired calcium images from the camera at a frame rate of 24 Hz. The software also triggered the vertical displacement of the light-sheet and the piezo nanopositioner, synchronized with the exposure of each frame, via a parallel port. This optical configuration generated a 24-layer stack with a temporal resolution of 1 Hz.

### Behavior recording and calcium imaging of two-photon microscopy

The fish was illuminated from the bottom (in behavioral tests) and above (near the objective lens, in the two-photon imaging experiments) by high power infrared light-emitting diodes (740 nm wavelength). To avoid photon interference, the experiments were carried out in the dark. The eye and tail movements were imaged with a resolution of 320 × 240 pixels at 200 frames per second using a CCD camera (PDV, MVC3000F-S00) mounted with a band-pass optical filter (central frequency: 740nm, FWHM: 40nm). The infrared light was reflected by a low pass filter (620 nm low-pass) on the light path of the visual stimuli. During the calcium imaging experiment, two-photon (Olympus, FV1000) laser (Mai Tai) was turned to 910 nm for excitation. Single slice of fish larval hindbrain was acquired every 1.1 seconds with a resolution of 512 × 512 pixels, during which the visual stimulus was presented and the behavioral responses were recorded. A 20x objective lens (Olympus, N20X-PFH) was used and the field-of-view is 318 × 318µm^2^, resulting in a spatial resolution of 0.62 × 0.62µm^2^ for each pixel. For each run of the test, 230 images (512 × 512) were collected.

### Behavior recording and calcium imaging of light-sheet microcopy

The behavioral responses of the fish were recorded at 200 frames per second using a CCD camera (PDV, MV-500C) with a resolution of 320 × 240 pixels. Infrared light from 6 high power LED (740 nm) was coupled into the light path of the imaging system to illuminate the head of the fish. In addition, the light from an infrared LED illuminated the tail of the fish via an optic fiber attached to the objective lens. The laser beam was projected onto the hindbrain from the left side of the fish creating the light-sheet which was moved in steps of 5µm, thereby covering most of the hindbrain (Ahrens et al., 2013; Panier et al., 2013). To minimize laser interference to the eye, a plastic opaque shutter was inserted in the agarose near the eye (Figure S4_1B). For each run of the experiment, 69 volumes of the images (2560 × 2160 × 24) were acquired.

### Behavior tracking and analysis

The behavioral image in RAW format was converted to 320 × 240 pixels PNG files by Matlab (MathWork) script. ImageJ (ImageJ 1.48v, NIH, USA) plugin Analyze Particles was used to track eye movements. By applying a proper threshold on the image brightness, eye was first segmented from the background and then fitted as an ellipse. The orientation of the longer axis of ellipse was used as the index for eye position. To determine the tail position, a quadrilateral was defined manually in a customized Matlab software. The quadrilateral usually was close to the tip of the tail and covered the part of the tail with best contrast from the background. The center of mass was calculated after the tail was segmented from the background. Horizontal positions were normalized in relative to the width of the quadrilateral, generating the index of tail position *m(t)* in each run of the experiments respectively. In the behavioral tests, the curve of eye positions was smoothed by a moving window averaging of 15 frames before the eye velocity was calculated as the differential of eye positions between successive time points. Tail beats were detected with a threshold of 0.2 and only one tail beat was allowed in every 2 s. The curves of eye velocity within the temporal window of 0.7 s before and 1s after the tail beats were collected and sorted by the stimulus status (moving or static). The tail beat induced responses on eye velocity were then averaged for moving and static stimuli separately. Paired *t-test* was used to compare the differences of the amplitude and latency of the peaks. For two-photon imaging and light-sheet imaging experiments, the curve of eye positions was resampled to 20 Hz before the velocity was calculated. The curve of eye positions was segmented into saccades and smooth pursuit (slow phase) according to the velocity by a threshold of 99% of the velocity distribution histogram. The averaged velocities of the smooth pursuit were compared between tail-immobilized runs and tail-free runs by paired *t-test*. The curve of tail position was also resampled to 20 Hz, before convolution with GC5 (GCaMP5g decay time constant) kernel (Kubo et al., 2014; Portugues et al., 2014) to generate the functional regressor (as described below). Tail beats were also detected by a threshold of 0.0625 and were used to categorize saccades into either TBIS, if a tail beat was detected within time frame of 250 ms before saccade onset, or normal saccade (no tail beat, Figure 6A). The TBISs were further divided into two sub-categories: saccades generated during moving gratings (Moving grating condition) and saccades generated during static gratings (Static grating condition).

### Analysis of two-photon images

The original images were converted to PNG format by FV1000 Viewer (Olympus) and were aligned by the TurboReg plugin in ImageJ to compensate for xy plane drifts (Sumbre et al., 2008). For each run, an anatomical reference was generated by averaging the realigned images. To measure the functional roles of the zebrafish hindbrain, eye position of each eye was convolved with an exponential kernel (the GCaMP5G decay time constant, GC5 kernel) to generate the OKR regressors, and a tail regressor was defined in the same manner (Kubo et al., 2014; Portugues et al., 2014). The co-registered images were filtered with a Gaussian low pass filter in spatial domain (size 5, sigma 2) and the intensity changes of each pixel across time were calculated as normalized ΔF/F (calcium transient), while the baseline (F) was defined as a polynomial function of fourth degree fitted to the curve of the pixel and the changes (ΔF) were obtained by subtracting the baseline from the original curve (Sumbre et al., 2008). Correlation maps were calculated for each run and each regressor, by pair wise correlation between the normalized ΔF/F of each pixel and the regressors. To obtain region of interests (ROIs) corresponding to individual neurons independently, a measurement of ‘peaky-ness’ (Ahrens et al., 2012) was also applied on the aligned images. A 11 × 11 moving window had been applied to each pixel in this process. The value of ‘peaky-ness’ for a pixel was defined as:

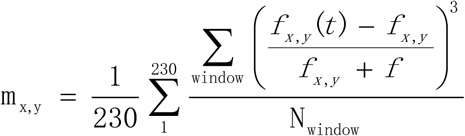

Where N_window_ is the number of valid pixels in the moving window centered at the position (x, y), f_x,y_(t) is the fluorescence of pixel (x,y) at time t (1 to 230), f_x,y_ is the average of fluorescence at pixel (x,y) and f is the average of f_x,y_ (across the imaging plane). The center of individual neurons were found for m_x,y_ with a peak larger than a preset threshold and at least 10 pixels from nearby peaks. For each correlation map, a valid neuron was defined as active pixels (r > 0.35) surrounding peak when they were neighboring to form a large enough (30 - 1200 pixels) connected cluster. For both tail conditions, the coordinates of individual neurons were manually matched between the 2 runs and only neurons with overlapping coordinates in both runs were selected as sensitive neurons. The layouts of sensitive neurons from all fish were merged for each condition while the intensity map represents the number of fish (count of the overlapping neurons, Figure 3C, 3D) or averaged correlation coefficient (Figure S3_1A). Three rectangular ROIs were defined in the merged maps (Figure 3C, 3D, Figure S3_1, Figure S3_2) and the differences of the tail-immobilized and tail-free conditions were tested separately for the three ROIs by paired *t-test*.

### Analysis of the light-sheet images

The calcium images from the light-sheet microscope were analyzed with Analysis of Functional NeuroImages suite (AFNI, NIH, (Cox, 1996)). The original image stacks of each run were converted to 3D+t (2592 × 2048 × 24 × 69) 3Df format and imported into AFNI. The dataset was registered to the first stack by a volume-based co-registration algorithm (3dvolreg, AFNI) to correct for minor motion artefacts in six axes (x, y, z, pitch, yaw, and roll). An anatomical reference was then generated for each run by averaging across the time domain. The anatomical references from the two runs were aligned by rigid body transformation (3dTagalign, AFNI) with the help of four markers defined in the images. The four markers were selected from neurons with high contrast to the background that existed in both runs, scattering in different depth and positions of the rostral-caudal axis of the hindbrain. These four markers, though variable across individual fish, were reliably recognized in different runs of the same fish. The aligned two datasets were then normalized to the Z-Brain Atlas template brain (Randlett et al., 2015) by an affine transformation, with the help of another four manually defined anatomical markers (Figure S4_3, one at the dorsal-caudal edge of the hindbrain rhombencephalon neuropil region 2, two at the most lateral edge close to left and right edge tip of rhombencephalon rhombomere 1, and one at the center of rhombencephalon rhombomere 3 below cerebellum, ZBrainViewer, ZBrain). This normalization procedure not only aligned the light-sheet datasets to the Z-Brain Atlas template brain, but also resampled the grids of images into 622 × 1406 × 138 with a resolution of 1 × 1 × 2 μm. The normalized datasets were converted to percent-signal-change (PSC or ΔF/F) for each voxel. First, system variance (defined by the signals outside of a brain mask drawn for each fish manually based on its anatomical reference) was removed from the raw fluorescence signals with 2nd order linear fit. Then the remaining curve was converted into ΔF/F timeseries similar as the procedure in two-photon imaging analysis (Figure 4A, also see SMovie4 and SMovie5). There were three regressors defined: the OKR regressor from the eye positions, a stimulus position regressor, and a saccade regressor. The OKR and stimulus position regressors were generated in the same manner as in the two-photon experiments. To generate the saccade regressor, the saccades detected in the curve of eye positions were replaced by a pulse function with duration of 1, while other data points of the curve were set to 0, and the resulting binary curve was convolved with the GC5 kernel (Figure S6). Functional activation maps were calculated by measuring the maximum correlation coefficients between the calcium responses (PSC) and the regressors voxel-wise with the 3ddelay program (AFNI), which produces both the value of the maximum coefficient and the latency of it. A spatial Gaussian filter with a radius of 5 was applied on the coefficient maps. The coefficient maps were first tested by one sample *t-test* against 0 at the individual level and group level. Then the differences between tail-immobilized and tail-free conditions were tested by a paired *t-test* at group level. To correct for potential probability bias from multiple comparisons, the minimum size of an activated cluster was determined by Monte Carlo simulations (AlphaSim, AFNI) and small clusters were excluded from the functional results at a given statistical threshold.

### Autoregressive model and Granger causality

A Matlab signal processing toolbox with the capability of vector autoregressive modeling, Automatic Spectral Analysis (http://www.mathworks.com/matlabcentral), was used to estimate the parameters of the autoregressive model, auto- and cross-correlations and spectra. Autoregressive modeling was implemented for each pair of signals based on a two-run procedure. In the first run, parameters for each pair of signals were selected automatically using a cross-entropy information criterion (de Waele and Broersen, 2003). Results revealed that no single pair had a best order larger than 5 and the largest best order for the group of pairs was determined by the first run. In the second run, using the largest best order, estimation of the model parameters, autocorrelations and cross correlations, spectra and directional Granger causality spectra (described below) was calculated trial by trial, before averaged across trials as a final stable estimation for that pair of signals. Directional Granger causality spectra could be calculated based on autoregressive modeling. When the coherence spectrum is defined as

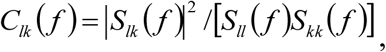

where S_lk_(f) is cross spectrum and S_ll_(f) and S_kk_(f) are auto-spectrum, the Granger causality spectrum from sensor l to k can be defined as

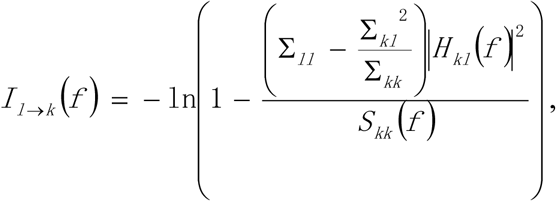

where Σ_ll_, Σ_kl_ and Σ_kk_ are elements of Σ, the covariance matrix, and S_kk_(f) is the power spectrum of sensor k. As a previous study had demonstrated (Brovelli et al., 2005), the frequency decomposition of the Granger causality could be interpreted as the amount of variance of signals from sensor k explained by the history of signals from sensor l.

To explore the informational contribution of the ROIs found in the comparison between tail-immobilized and tail-free conditions in the light-sheet imaging experiment (Figure 3D), the Granger causality was calculated for the rostral hindbrain (rHB) ROI and dorsal-caudal hindbrain (dcHB) ROI separately. The normalized datasets (622 × 1406 × 138) was divided into 4 × 4 × 4 bricks. The information flow was measured at brick level between the voxels in rHB/dcHB and all the bricks. For each pair of ROI-bricks, there are Num_roi_voxel_ × 64 combinations of signals, in which the Num_roi_voxel_ is the number of voxels in a given ROI (rHB or dcHB) and each brick has 64 voxels. The Granger causality was calculated for each bricks as if there were Num_roi_voxel_ × 64 repetition of trials, as described above. The Granger values, indicating information to and from the ROI, were filled into corresponding 64 voxels to generate the information flow maps, separately. A spatial Gaussian filter with a radius of 5 was applied on the information flow maps. The resulting maps were tested at group level by one-sample *t-test* against 0. The same clustering parameters for multiple correction were applied as that used in the functional analysis.

### Analysis of saccade induced neural level responses

The calcium responses of each sensitive neuron in the two-photon experiment were extracted and split into short episodes (11 data points, −3.3s to 7.7s, relative to the onset of the saccades). The baseline drift of each episode was removed by subtracting the value of the first data point from all the 11 data points. Saccade-induced response of each episode was evaluated by the difference between two intervals around the saccade (−3.3 −0s vs. 1.1 −4.4s). The episodes with mean difference larger than 3 (ΔF/F) were selected and sorted into three groups by the categories of the saccades (Moving grating, static grating, and no tail beat). Averaged traces of the three groups of episodes were compared for each time point pair-wise (Figure 6A). Within each group, the episodes were sorted by the neural response at the saccade (Figure 6B) before the averaging of the first half of the episodes was compared with the averaging of the second half of the episodes (Figure 6C).

## References

Ahrens, M.B., Li, J.M., Orger, M.B., Robson, D.N., Schier, A.F., Engert, F., and Portugues, R. (2012). Brain-wide neuronal dynamics during motor adaptation in zebrafish. Nature 485, 471–477.

Ahrens, M.B., Orger, M.B., Robson, D.N., Li, J.M., and Keller, P.J. (2013). Whole-brain functional imaging at cellular resolution using light-sheet microscopy. Nat Methods 10, 413–420.

Ambati, V.N., Saucedo, F., Murray, N.G., Powell, D.W., and Reed-Jones, R.J. (2016). Constraining eye movement in individuals with Parkinson’s disease during walking turns. Exp Brain Res 234, 2957–2965.

Andreescu, C.E., De Ruiter, M.M., De Zeeuw, C.I., and De Jeu, M.T. (2005). Otolith deprivation induces optokinetic compensation. J Neurophysiol 94, 3487–3496.

Angelaki, D.E., and Hess, B.J. (2005). Self-motion-induced eye movements: effects on visual acuity and navigation. Nat Rev Neurosci 6, 966–976.

Arrenberg, A.B., Del Bene, F., and Baier, H. (2009). Optical control of zebrafish behavior with halorhodopsin. Proc Natl Acad Sci U S A 106, 17968–17973.

Barker, A.J., and Baier, H. (2015). Sensorimotor Decision Making in the Zebrafish Tectum. Current Biology 25, 2804–2814.

Bell, C.C. (1981). An efference copy which is modified by reafferent input. Science 214, 450–453.

Berkes, P., Orban, G., Lengyel, M., and Fiser, J. (2011). Spontaneous Cortical Activity Reveals Hallmarks of an Optimal Internal Model of the Environment. Science 331, 83–87.

Bhattacharyya, K., McLean, D.L., and MacIver, M.A. (2017). Visual Threat Assessment and Reticulospinal Encoding of Calibrated Responses in Larval Zebrafish. Curr Biol 27, 2751–2762 e2756.

Bianco, I.H., and Engert, F. (2015). Visuomotor transformations underlying hunting behavior in zebrafish. Curr Biol 25, 831–846.

Braitenberg, V., Heck, D., and Sultan, F. (1997). The detection and generation of sequences as a key to cerebellar function: experiments and theory. The Behavioral and brain sciences 20, 229–245; discussion 245-277.

Brovelli, A., Lachaux, J.P., Kahane, P., and Boussaoud, D. (2005). High gamma frequency oscillatory activity dissociates attention from intention in the human premotor cortex. Neuroimage 28, 154– 164.

Chagnaud, B.P., Simmers, J., and Straka, H. (2012). Predictability of visual perturbation during locomotion: implications for corrective efference copy signaling. Biol Cybern 106, 669–679.

Combes, D., Le Ray, D., Lambert, F.M., Simmers, J., and Straka, H. (2008). An intrinsic feed-forward mechanism for vertebrate gaze stabilization. Curr Biol 18, R241–243.

Cox, R.W. (1996). AFNI: software for analysis and visualization of functional magnetic resonance neuroimages. Computers and biomedical research, an international journal 29, 162–173.

Cullen, K.E. (2004). Sensory signals during active versus passive movement. Curr Opin Neurobiol 14, 698–706.

Daie, K., Goldman, M.S., and Aksay, E.R. (2015). Spatial patterns of persistent neural activity vary with the behavioral context of short-term memory. Neuron 85, 847–860.

de Waele, S., and Broersen, P.M.T. (2003). Order selection for vector autoregressive models. Ieee T Signal Proces 51, 427–433.

Del Bene, F., Wyart, C., Robles, E., Tran, A., Looger, L., Scott, E.K., Isacoff, E.Y., and Baier, H. (2010). Filtering of visual information in the tectum by an identified neural circuit. Science 330, 669–673.

Dunn, T.W., Gebhardt, C., Naumann, E.A., Riegler, C., Ahrens, M.B., Engert, F., and Del Bene, F. (2016). Neural Circuits Underlying Visually Evoked Escapes in Larval Zebrafish. Neuron 89, 613– 628.

Easter, S.S., and Johns, P.R. (1974). Horizontal Compensatory Eye-Movements in Goldfish (Carassius-Auratus). 2. Comparison of Normal and Deafferented Animals. J Comp Physiol 92, 37– 57.

Easter, S.S., and Nicola, G.N. (1997). The development of eye movements in the zebrafish (Danio rerio). Dev Psychobiol 31, 267–276.

Edwards, D.H., Heitler, W.J., and Krasne, F.B. (1999). Fifty years of a command neuron: the neurobiology of escape behavior in the crayfish. Trends in neurosciences 22, 153–161.

Fallani Fde, V., Corazzol, M., Sternberg, J.R., Wyart, C., and Chavez, M. (2015). Hierarchy of neural organization in the embryonic spinal cord: Granger-causality graph analysis of in vivo calcium imaging data. IEEE Trans Neural Syst Rehabil Eng 23, 333–341.

Frens, M.A., and Donchin, O. (2009). Forward models and state estimation in compensatory eye movements. Frontiers in cellular neuroscience 3, 13.

Gibson, J.J. (1972). A theory of direct visual perception. In Vision and Mind: Selected Readings in the Philosophy of Perception, A. Noe, and E. Thompson, eds. (MIT Press), pp. 77–89.

Godaux, E., and Vanderkelen, B. (1984). Vestibulo-ocular reflex, optokinetic response and their interactions in the cerebellectomized cat. J Physiol 346, 155–170.

Goncalves, P.J., Arrenberg, A.B., Hablitzel, B., Baier, H., and Machens, C.K. (2014). Optogenetic perturbations reveal the dynamics of an oculomotor integrator. Frontiers in neural circuits 8, 10.

Granger, C.W.J. (1969). Investigating Causal Relations by Econometric Models and Cross-Spectral Methods. Econometrica 37, 424–438.

Green, A.M., and Angelaki, D.E. (2010). Multisensory integration: resolving sensory ambiguities to build novel representations. Curr Opin Neurobiol 20, 353–360.

Gregoriou, G.G., Gotts, S.J., Zhou, H., and Desimone, R. (2009). High-frequency, long-range coupling between prefrontal and visual cortex during attention. Science 324, 1207–1210.

Grillner, S., Ekeberg, El Manira, A., Lansner, A., Parker, D., Tegner, J., and Wallen, P. (1998). Intrinsic function of a neuronal network - a vertebrate central pattern generator. Brain research Brain research reviews 26, 184–197.

Huang, Y.Y., and Neuhauss, S.C. (2008). The optokinetic response in zebrafish and its applications. Frontiers in bioscience: a journal and virtual library 13, 1899–1916.

Kaneko, M., Hernandez-Borsetti, N., and Cahill, G.M. (2006). Diversity of zebrafish peripheral oscillators revealed by luciferase reporting. Proc Natl Acad Sci U S A 103, 14614–14619.

Kennedy, D., Calabrese, R.L., and Wine, J.J. (1974). Presynaptic inhibition: primary afferent depolarization in crayfish neurons. Science 186, 451–454.

Kilner, J.M., Friston, K.J., and Frith, C.D. (2007). Predictive coding: an account of the mirror neuron system. Cognitive processing 8, 159–166.

Knight, T.A. (2012). Contribution of the frontal eye field to gaze shifts in the head-unrestrained rhesus monkey: neuronal activity. Neuroscience 225, 213–236.

Knogler, L.D., Markov, D.A., Dragomir, E.I., Stih, V., and Portugues, R. (2017). Sensorimotor Representations in Cerebellar Granule Cells in Larval Zebrafish Are Dense, Spatially Organized, and Non-temporally Patterned. Curr Biol 27, 1288–1302.

Koyama, M., Kinkhabwala, A., Satou, C., Higashijima, S., and Fetcho, J. (2011). Mapping a sensorymotor network onto a structural and functional ground plan in the hindbrain. Proceedings of the National Academy of Sciences of the United States of America 108, 1170–1175.

Kubo, F., Hablitzel, B., Dal Maschio, M., Driever, W., Baier, H., and Arrenberg, A.B. (2014). Functional architecture of an optic flow-responsive area that drives horizontal eye movements in zebrafish. Neuron 81, 1344–1359.

Lambert, F.M., Combes, D., Simmers, J., and Straka, H. (2012). Gaze stabilization by efference copy signaling without sensory feedback during vertebrate locomotion. Curr Biol 22, 1649–1658.

Lisberger, S.G. (2009). Internal models of eye movement in the floccular complex of the monkey cerebellum. Neuroscience 162, 763–776.

Llinas, R.R. (1988). The intrinsic electrophysiological properties of mammalian neurons: insights into central nervous system function. Science 242, 1654–1664.

Miri, A., Daie, K., Arrenberg, A.B., Baier, H., Aksay, E., and Tank, D.W. (2011). Spatial gradients and multidimensional dynamics in a neural integrator circuit. Nat Neurosci 14, 1150–1159.

Mu, Y., Li, X.Q., Zhang, B., and Du, J.L. (2012). Visual Input Modulates Audiomotor Function via Hypothalamic Dopaminergic Neurons through a Cooperative Mechanism. Neuron 75, 688–699.

Neuhauss, S.C.F. (2003). Behavioral genetic approaches to visual system development and function in zebrafish. J Neurobiol 54, 148–160.

Panier, T., Romano, S.A., Olive, R., Pietri, T., Sumbre, G., Candelier, R., and Debregeas, G. (2013). Fast functional imaging of multiple brain regions in intact zebrafish larvae using selective plane illumination microscopy. Frontiers in neural circuits 7, 65.

Perez-Schuster, V., Kulkarni, A., Nouvian, M., Romano, S.A., Lygdas, K., Jouary, A., Dipoppa, M., Pietri, T., Haudrechy, M., Candat, V., et al. (2016). Sustained Rhythmic Brain Activity Underlies Visual Motion Perception in Zebrafish. Cell reports 17, 3089.

Portugues, R., Feierstein, C.E., Engert, F., and Orger, M.B. (2014). Whole-brain activity maps reveal stereotyped, distributed networks for visuomotor behavior. Neuron 81, 1328–1343.

Randlett, O., Wee, C.L., Naumann, E.A., Nnaemeka, O., Schoppik, D., Fitzgerald, J.E., Portugues, R., Lacoste, A.M.B., Riegler, C., Engert, F., and Schier, A.F. (2015). Whole-brain activity mapping onto a zebrafish brain atlas. Nature Methods 12, 1039–1046.

Renninger, S.L., Schonthaler, H.B., Neuhauss, S.C.F., and Dahm, R. (2011). Investigating the genetics of visual processing, function and behaviour in zebrafish. Neurogenetics 12, 97–116.

Roebroeck, A., Formisano, E., and Goebel, R. (2005). Mapping directed influence over the brain using Granger causality and fMRI. Neuroimage 25, 230–242.

Romano, S.A., Pietri, T., Perez-Schuster, V., Jouary, A., Haudrechy, M., and Sumbre, G. (2015). Spontaneous neuronal network dynamics reveal circuit’s functional adaptations for behavior. Neuron 85, 1070–1085.

Rosch, R.E., Hunter, P.R., Baldeweg, T., Friston, K.J., and Meyer, M.P. (2018). Calcium imaging and dynamic causal modelling reveal brain-wide changes in effective connectivity and synaptic dynamics during epileptic seizures. PLoS Comput Biol 14, e1006375.

Rosenberg, A.F., and Ariel, M. (1996). A model for optokinetic eye movements in turtles that incorporates properties of retinal-slip neurons. Vis Neurosci 13, 375–383.

Ross, J., Morrone, M.C., and Burr, D.C. (1997). Compression of visual space before saccades. Nature 386, 598–601.

Roy, J.E., and Cullen, K.E. (2001). Selective processing of vestibular reafference during selfgenerated head motion. J Neurosci 21, 2131–2142.

Roy, J.E., and Cullen, K.E. (2004). Dissociating self-generated from passively applied head motion: neural mechanisms in the vestibular nuclei. J Neurosci 24, 2102–2111.

Sadtler, P.T., Quick, K.M., Golub, M.D., Chase, S.M., Ryu, S.I., Tyler-Kabara, E.C., Yu, B.M., and Batista, A.P. (2014). Neural constraints on learning. Nature 512, 423–426.

Schoonheim, P.J., Arrenberg, A.B., Del Bene, F., and Baier, H. (2010). Optogenetic localization and genetic perturbation of saccade-generating neurons in zebrafish. J Neurosci 30, 7111–7120.

Semmelhack, J.L., Donovan, J.C., Thiele, T.R., Kuehn, E., Laurell, E., and Baier, H. (2014). A dedicated visual pathway for prey detection in larval zebrafish. eLife 3.

Shadmehr, R., Smith, M.A., and Krakauer, J.W. (2010). Error correction, sensory prediction, and adaptation in motor control. Annu Rev Neurosci 33, 89–108.

Shanidze, N., Kim, A.H., Loewenstein, S., Raphael, Y., and King, W.M. (2010a). Eye-head coordination in the guinea pig II. Responses to self-generated (voluntary) head movements. Exp Brain Res 205, 445–454.

Shanidze, N., Kim, A.H., Raphael, Y., and King, W.M. (2010b). Eye-head coordination in the guinea pig I. Responses to passive whole-body rotations. Exp Brain Res 205, 395–404.

Sommer, M.A., and Wurtz, R.H. (2002). A pathway in primate brain for internal monitoring of movements. Science 296, 1480–1482.

Sommer, M.A., and Wurtz, R.H. (2008). Brain circuits for the internal monitoring of movements. Annu Rev Neurosci 31, 317–338.

Sperry, R.W. (1950). Neural Basis of the Spontaneous Optokinetic Response Produced by Visual Inversion. J Comp Physiol Psych 43, 482–489.

Srivastava, A., Ahmad, O.F., Pacia, C.P., Hallett, M., and Lungu, C. (2018). The Relationship between Saccades and Locomotion. Journal of movement disorders 11, 93–106.

Stehouwer, D.J. (1987). Compensatory eye movements produced during fictive swimming of a deafferented, reduced preparation in vitro. Brain Res 410, 264–268.

Sumbre, G., Muto, A., Baier, H., and Poo, M.M. (2008). Entrained rhythmic activities of neuronal ensembles as perceptual memory of time interval. Nature 456, 102–106.

Sun, L.D., and Goldberg, M.E. (2016). Corollary Discharge and Oculomotor Proprioception: Cortical Mechanisms for Spatially Accurate Vision. Annual review of vision science 2, 61–84.

Temizer, I., Donovan, J.C., Baier, H., and Semmelhack, J.L. (2015). A Visual Pathway for Looming-Evoked Escape in Larval Zebrafish. Curr Biol 25, 1823–1834.

von Holst E, M.H. (1950). Das reafferenzprinzip. Naturwissenschaften 37, 464–476.

Warp, E., Agarwal, G., Wyart, C., Friedmann, D., Oldfield, C.S., Conner, A., Del Bene, F., Arrenberg, A.B., Baier, H., and Isacoff, E.Y. (2012). Emergence of patterned activity in the developing zebrafish spinal cord. Curr Biol 22, 93–102.

Winter, R., Harrar, V., Gozdzik, M., and Harris, L.R. (2008). The relative timing of active and passive touch. Brain Res 1242, 54–58.

Wolf, S., Dubreuil, A.M., Bertoni, T., Bohm, U.L., Bormuth, V., Candelier, R., Karpenko, S., Hildebrand, D.G.C., Bianco, I.H., Monasson, R., and Debregeas, G. (2017). Sensorimotor computation underlying phototaxis in zebrafish. Nature communications 8, 651.

Wolpert, D.M., and Miall, R.C. (1996). Forward Models for Physiological Motor Control. Neural Netw 9, 1265–1279.

Wurtz, R.H. (2018). Corollary Discharge Contributions to Perceptual Continuity Across Saccades. Annual review of vision science 4, 215–237.

Wyart, C., Del Bene, F., Warp, E., Scott, E.K., Trauner, D., Baier, H., and Isacoff, E.Y. (2009). Optogenetic dissection of a behavioural module in the vertebrate spinal cord. Nature 461, 407– 410.

Yao, Y.Y., Li, X.Q., Zhang, B.B., Yin, C., Liu, Y.F., Chen, W.Y., Zeng, S.Q., and Du, J.L. (2016). Visual Cue-Discriminative Dopaminergic Control of Visuomotor Transformation and Behavior Selection. Neuron 89, 598–612.

Yoder, R.M., Clark, B.J., Brown, J.E., Lamia, M.V., Valerio, S., Shinder, M.E., and Taube, J.S. (2011). Both visual and idiothetic cues contribute to head direction cell stability during navigation along complex routes. J Neurophysiol 105, 2989–3001.

