## Supplementary Figures for "Neural circuits mediating visual stabilization during active motion in zebrafish"

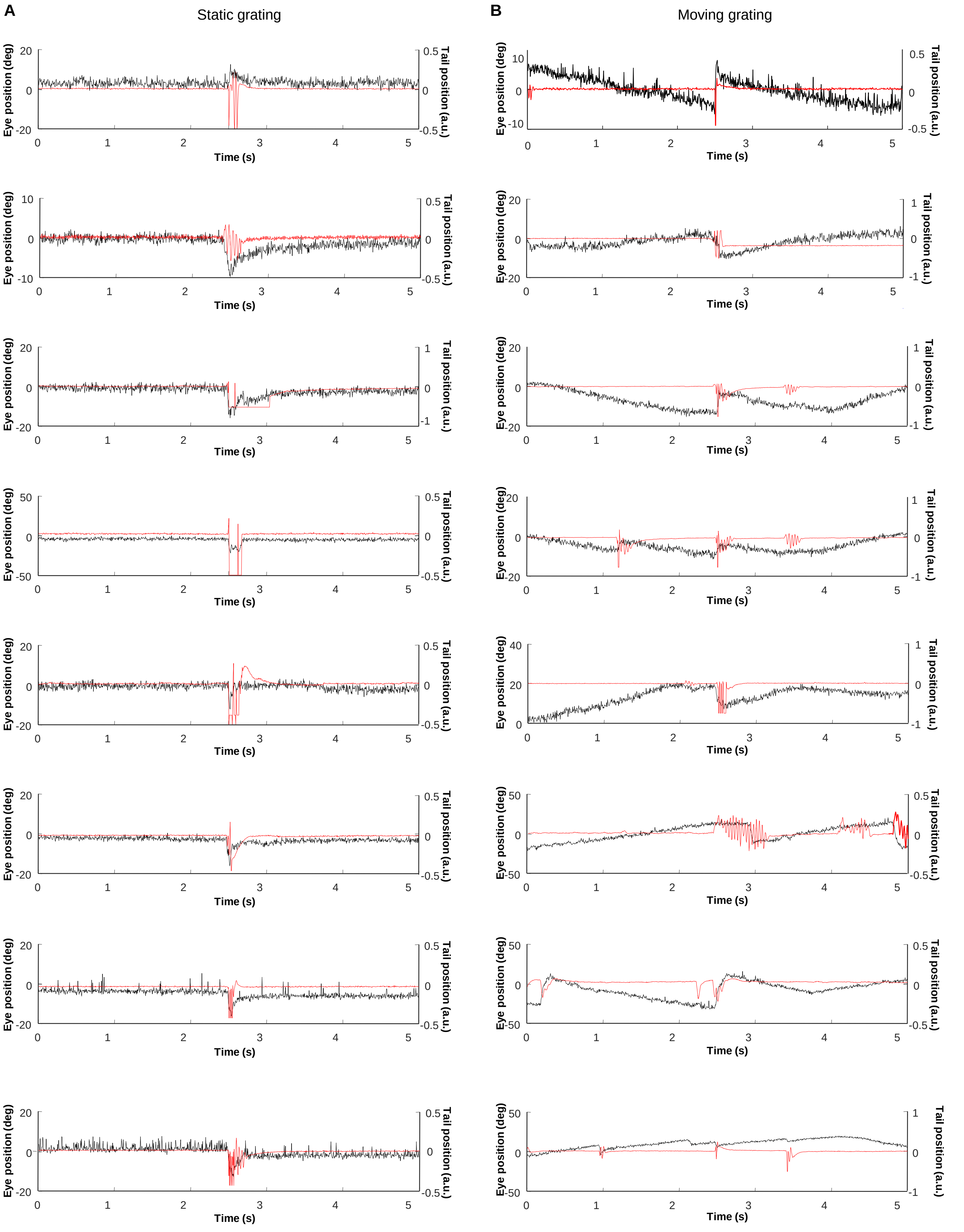


**Figure S1_1. More examples of tail-beat induced saccade.**
(**A**) Tail movements induced saccade during presentation of static grating. (**B**) Tail-beat resets eye position when presented with moving grating.

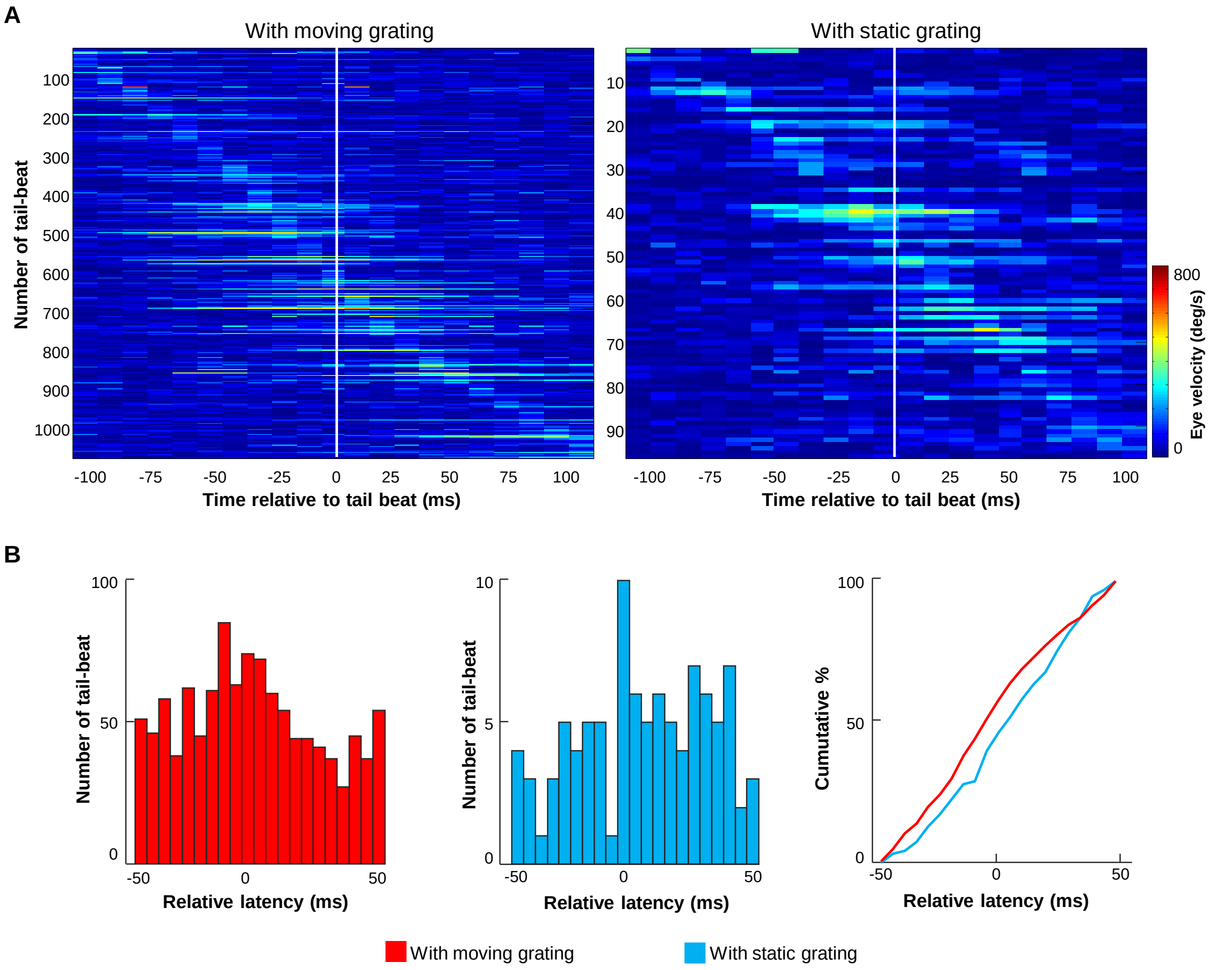


**Figure S1_2. Relative tail-saccade latency for moving/static grating.**
(**A**) Due to the design of the experiment, more tail beats were found during the moving grating runs. Eye velocity 50 ms before and 50 ms after a tail beat was shown, sorted by the latency of the peak. (**B**) The distribution of the relative tail-saccade latency for those tail beats is in favor of shorter latency during the moving grating (left panel), compared with those tail beats happened during the presention of the static grating (middle panel). The difference is significant ($P < 0.05$, right panel) by KS test.

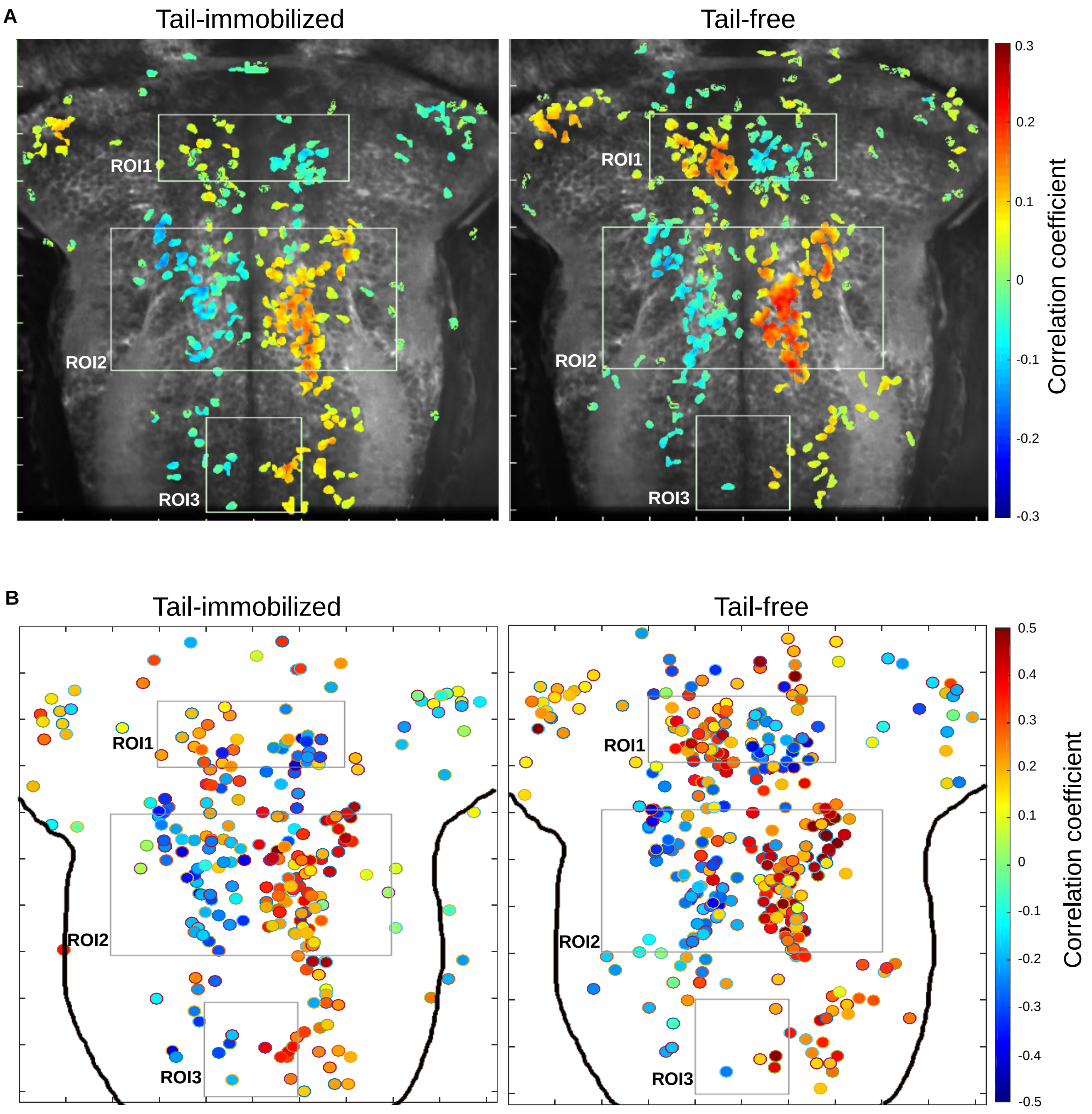


**Figure S3_1. Averaged response map and 2D reconstructed map of OKR responsive neurons.**

(**A**) OKR related neural responses in tail-immobilized condition (left) and tail-free condition (right) were pooled together across fish. Pseudocolor scale depicts the averaged correlation coefficient. (**B**) 2D reconstruction of the OKR responsive neurons. Each dot represents one neuron and is color-coded according to its correlation coefficient with the OKR regressor. The three rectangular boxes indicate the same ROIs as in Figure 2. Notice the inverted left-right polarity of the coefficient in ROI1 and ROI2/3.

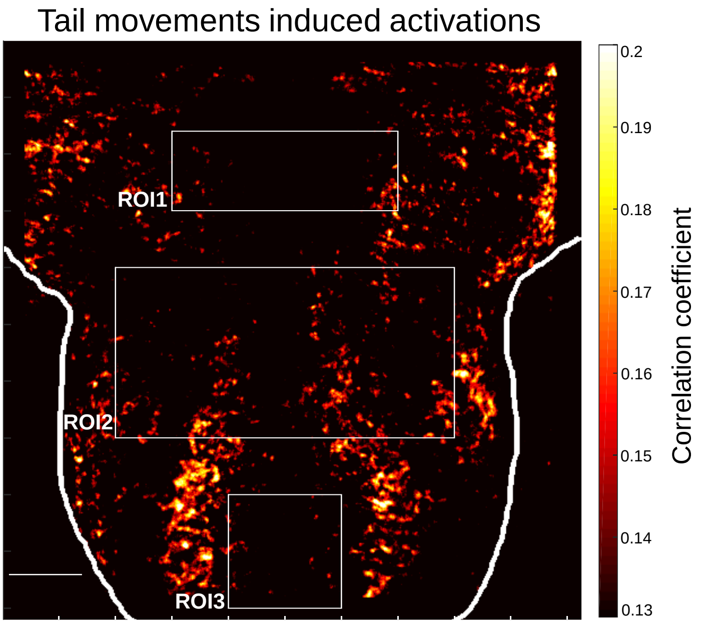


**Figure S3_2. Response map for tail movements.**
A tail movement regressor was generated in the same way as that of OKR regressor, based on which neural responses were calculated for correlation coefficient. The responses were averaged for tail-free condition across fish. Though there were some activations expanding rostrally into ROI2, most of the activations were found in caudal hindbrain, much lateral to ROI3.

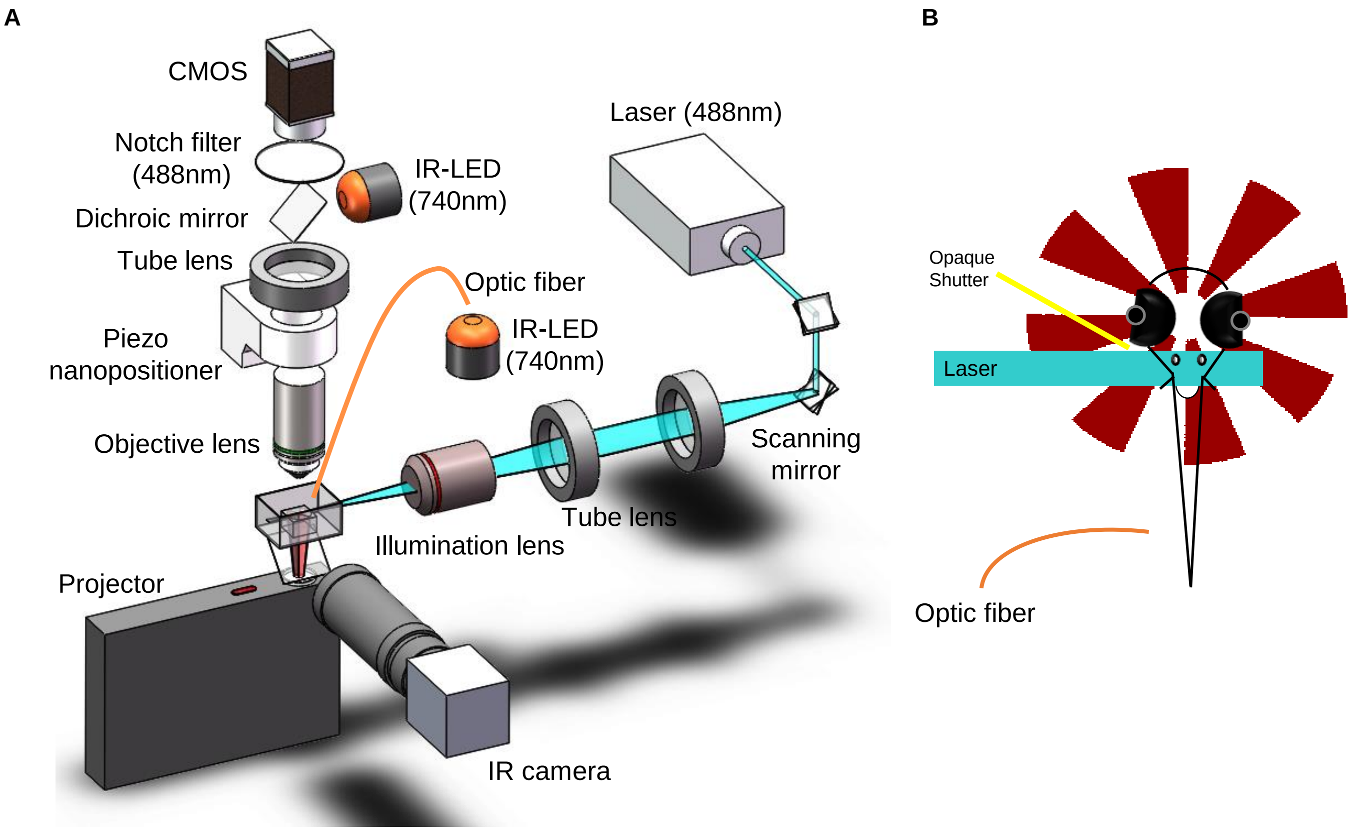


**Figure S4_1. Setup of volumetric imaging with light-sheet microscope.**
(**A**) Sketch of the optical set-up. In spite of embedding the visual presenting system into a custom light-sheet microscope, a dichroic mirror was mounted between the notch filter and the tube lens for the IR illumination on the eye, and an optic fiber was attached to the side of objective lens to highlight the tail. (**B**) Hindbrain was illuminated from side of the larvae, covering most depth of the hindbrain. With the help of a piece of opaque shutter, reliable OKR was observed even though the 488 nm laser for fluorescence excitation is visible.

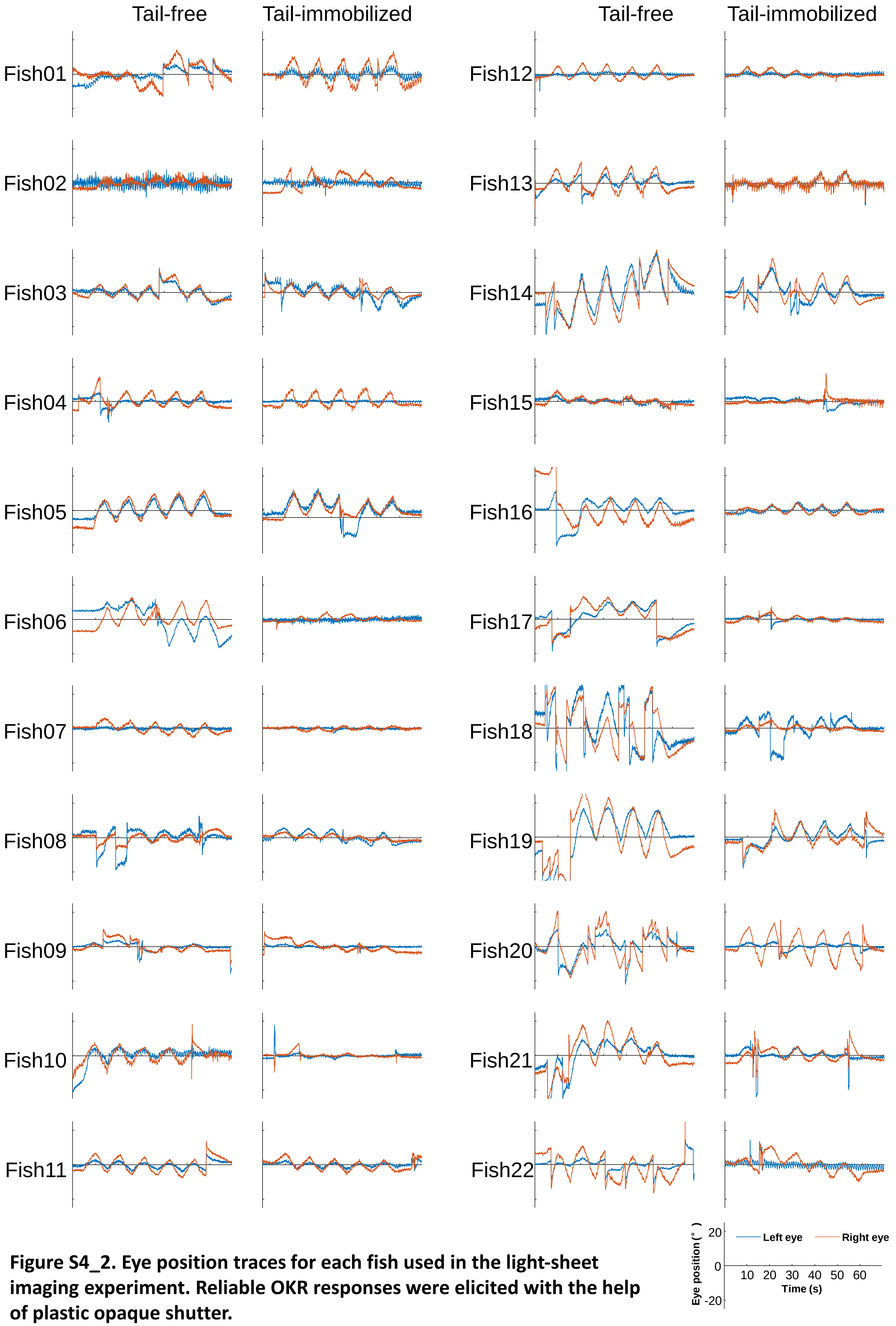


**Figure S4_2. Eye position traces for each fish used in the light-sheet imaging experiment. Reliable OKR responses were elicited with the help of plastic opaque shutter.**

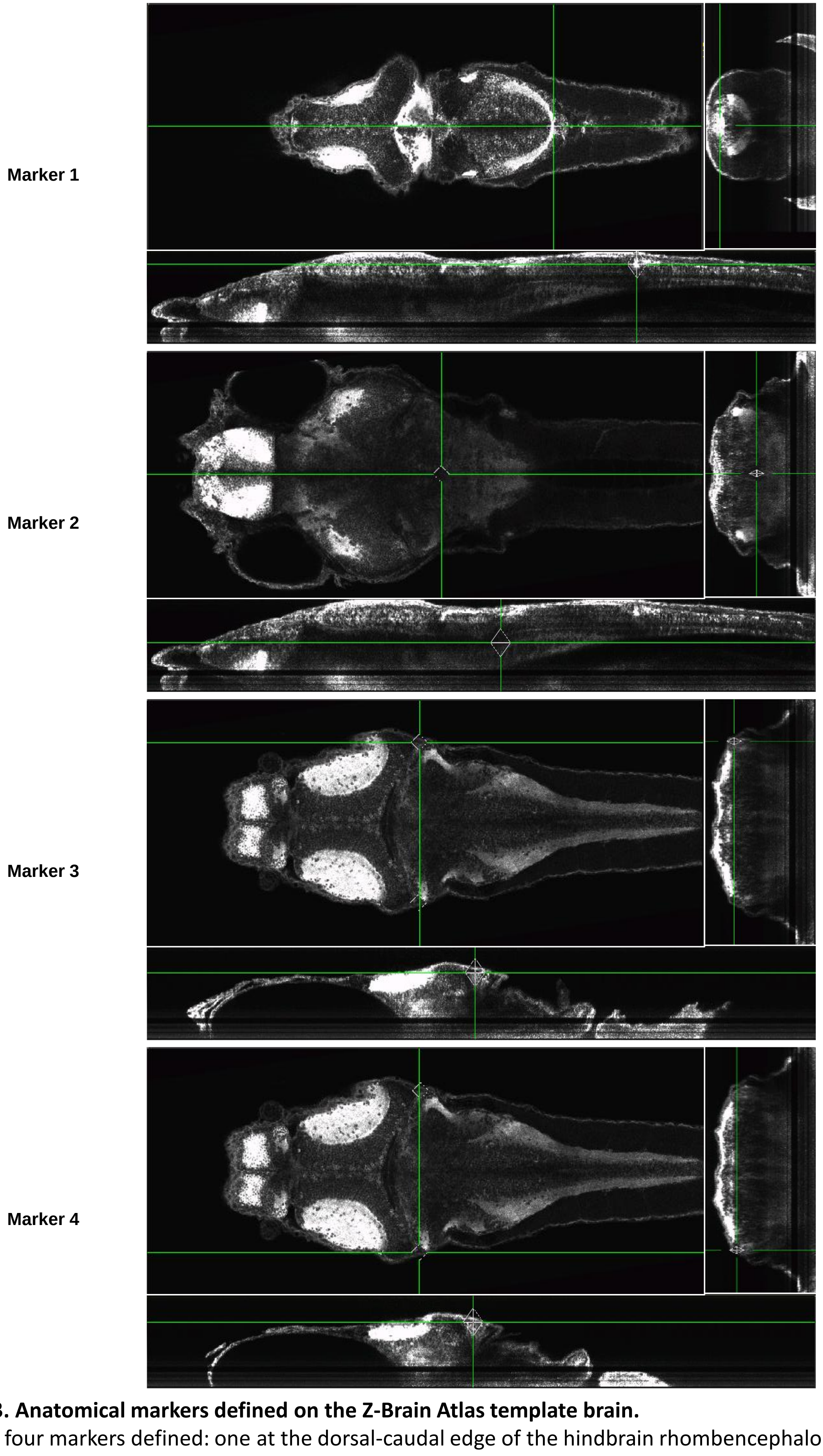


**Figure S4_3. Anatomical markers defined on the Z-Brain Atlas template brain.**
There were four markers defined: one at the dorsal-caudal edge of the hindbrain rhombencephalon neuropil region 2, two at the most lateral edge close to left and right edge tip of rhombencephalon rhombomere 1, and one at the center of rhombencephalon rhombomere 3 below cerebellum.

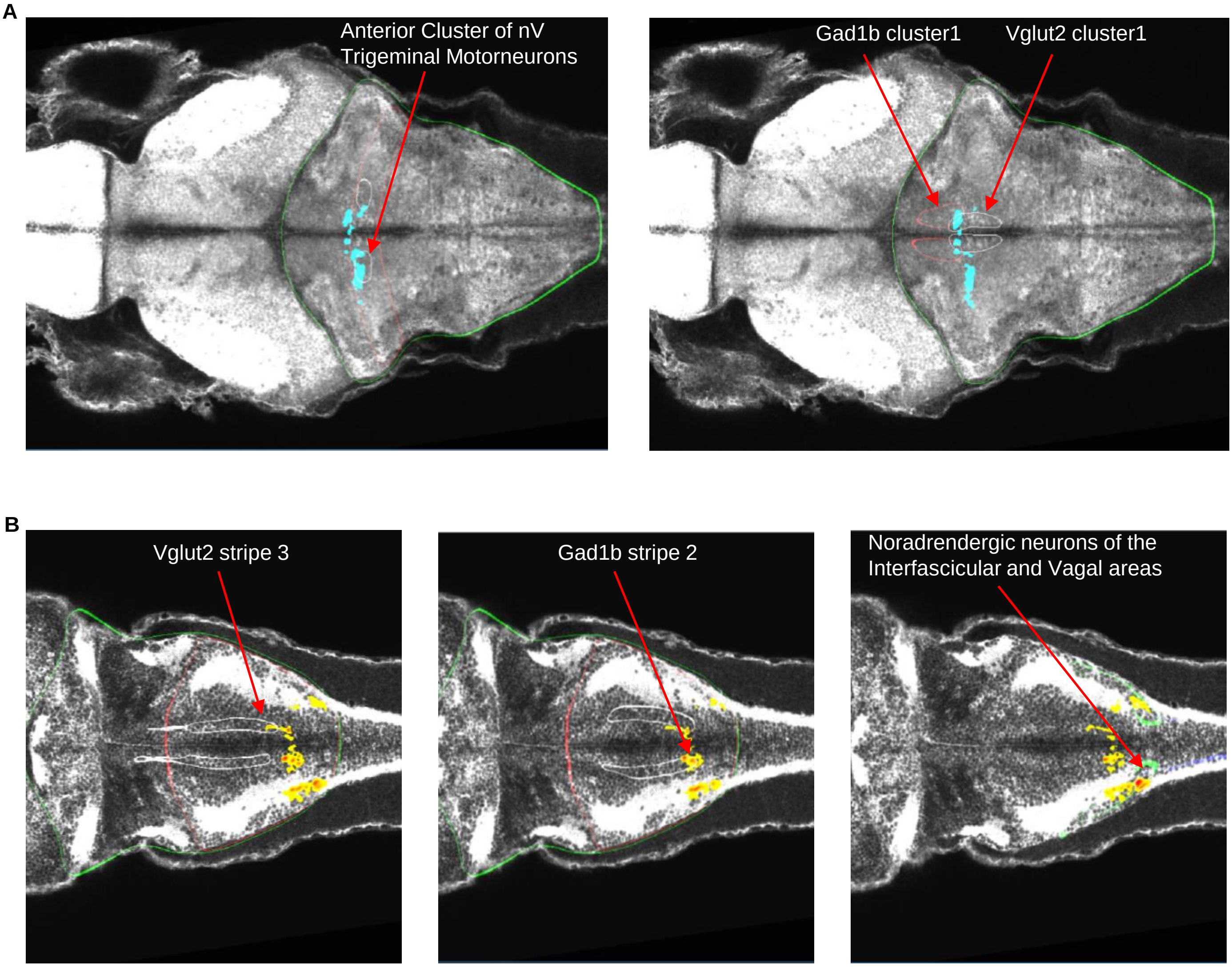


**Figure S4_4. Brain regions recognized in Z-Brain atlas.**
(**A**) For the activations larger for tail-free condition, the brain regions include anterior cluster of nV trigeminal motorneurons, Vglut2 Cluster 1 and Gad1b Cluster 1. (**B**) For tail-immobilized condition, the activations scattered among Gad1b Stripe 2, Vglut2 Stripe 3 and noradrendergic neurons of the interfascicular and Vagal areas.

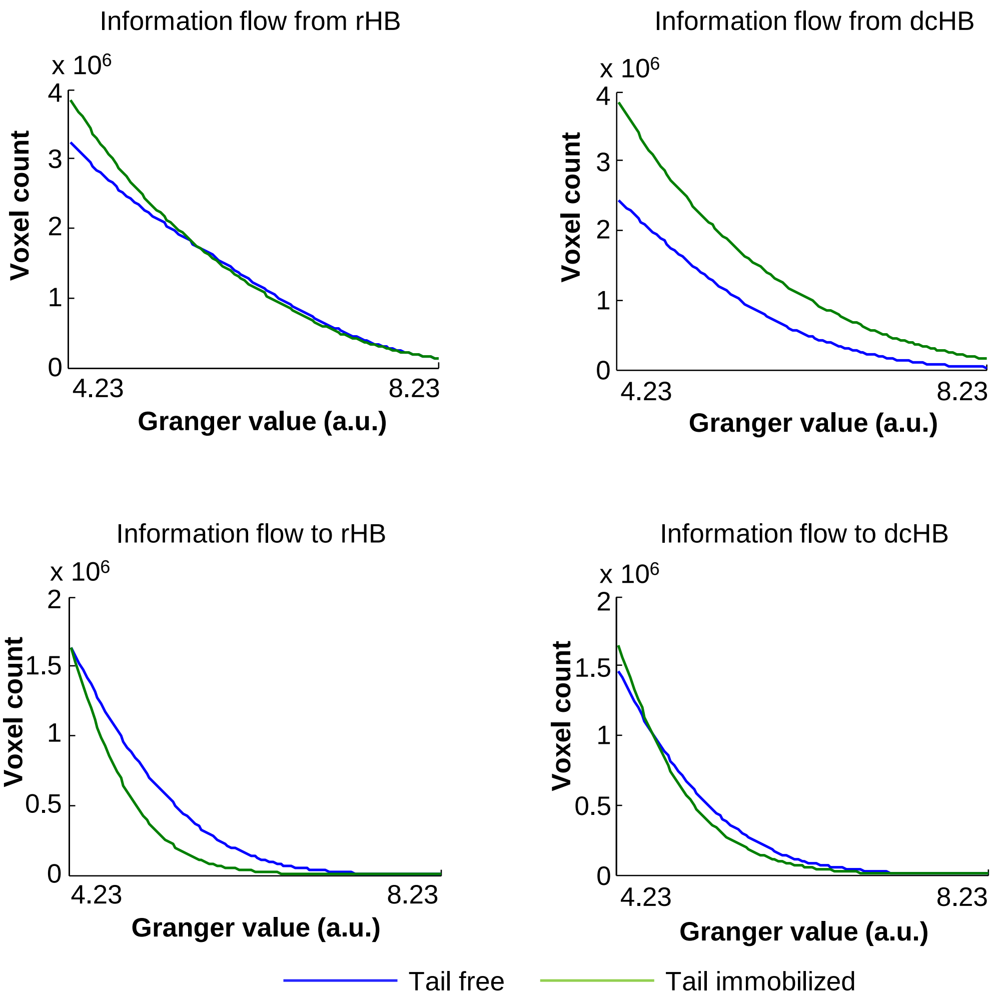


**Figure S5. Tail movements altered information flow in hindbrain.**
For the information flow into rHB, there were more voxels in hindbrain involved during tail-free condition. For tail-immobilized condition, there were more voxels in hindbrain receive information from dcHB. This pattern was consistent across a wide range of threshold. $P < 0.05$, into rHB; $P < 0.002$, from dcHB, KS-test.

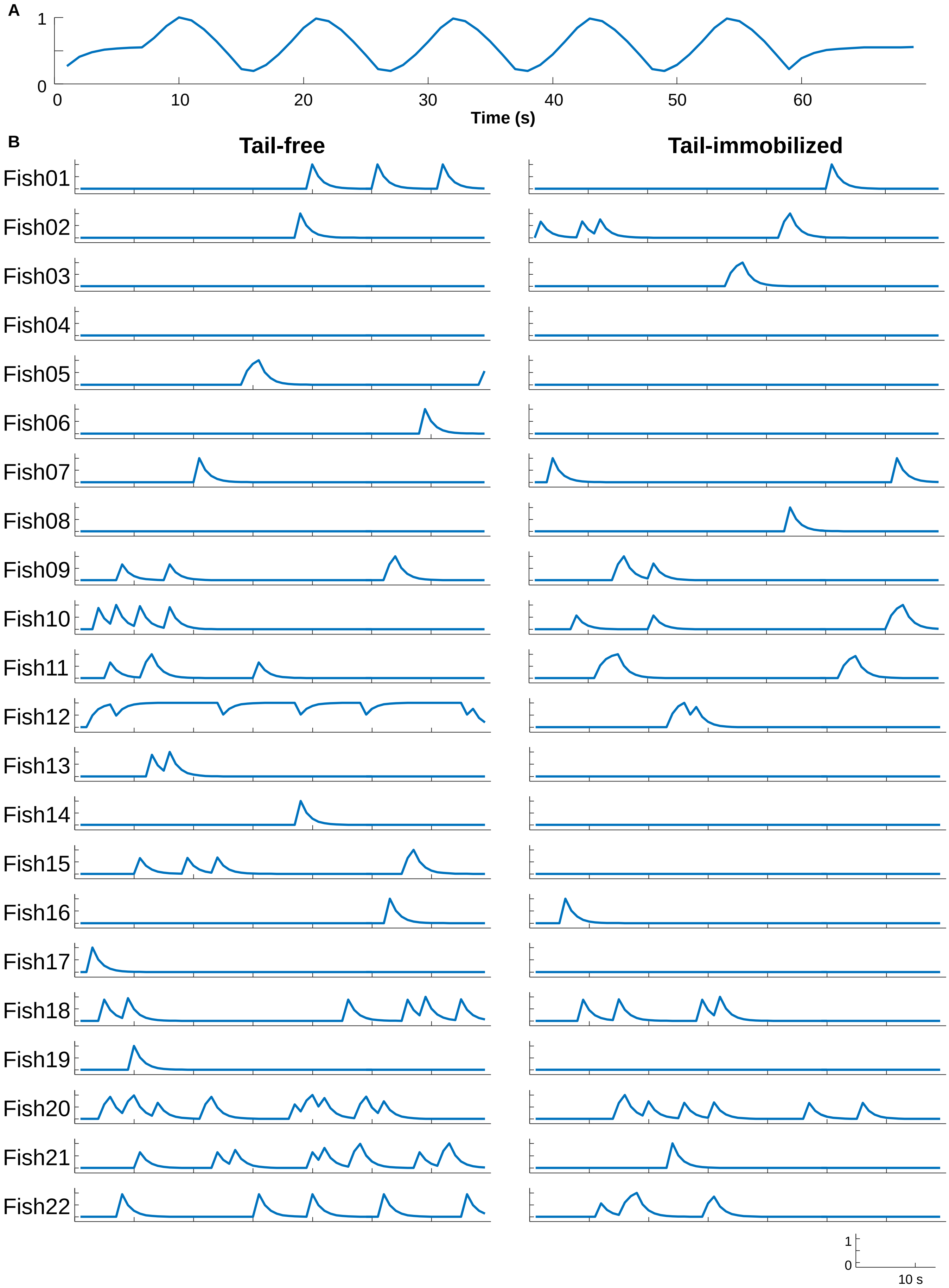


**Figure S6. Stimulus regressors and saccade regressors.**
(**A**) Stimulus position regressor for every fish in all conditions. (**B**) Saccade regressors for each fish in tail-free condition (left panel) and tail-immobilized condition (right panel).
